# A Generative Foundation Model for Cryo-EM Densities

**DOI:** 10.64898/2025.12.29.696802

**Authors:** Yilai Li, Jing Yuan, Yi Zhou, Zhenghua Wang, Suyi Chen, Fengyu Yang, Haibin Ling, Shahar Z. Kovalsky, Xiaoqing Zheng, Quanquan Gu

**Author notes:** Equal contribution (Yilai Li, Jing Yuan, Yi Zhou). Work done at ByteDance (Zhenghua Wang, Suyi Chen, Fengyu Yang).

## Abstract

Single-particle cryo-electron microscopy (cryo-EM) enables structure determination of macromolecular complexes, yet reconstructions are often degraded by noise, anisotropic sampling and reconstruction artifacts. Correcting these effects is an ill-posed inverse problem that requires strong prior knowledge in addition to experimental data. Here we introduce cryoFM, a generative foundation model for cryo-EM densities that unifies data-driven structural priors with dataset-specific constraints within a Bayesian inference framework. CryoFM is trained in an unsupervised manner on thousands of high-quality cryo-EM maps using flow matching, learning a generalizable prior over macromolecular density distributions. Combined with explicit likelihood models describing experimental degradations, cryoFM enables flow posterior sampling, an inference-only procedure that performs denoising and restoration and refinement while remaining explicitly constrained by dataset-derived statistics. We show that this framework improves density reconstruction and refinement across diverse experimental settings, including preferred-orientation datasets and cases with strong spatial heterogeneity in signal-to-noise ratio, without introducing hallucinated features. In addition, cryoFM can be fine-tuned into conditional generative models for density post-processing, yielding maps with improved interpretability and fewer artifacts compared to existing supervised approaches. Together, cryoFM establishes generative foundation models as a principled and controllable framework for cryo-EM density reconstruction and modification.

## Introduction

Single particle cryo-electron microscopy (cryo-EM) has rapidly become one of the most important tools in structural biology^1^. Despite its broad adoption, 3D reconstruction from single particle cryo-EM data continues to face many challenges. Small or flexible targets remain difficult, datasets often suffer from preferred particle orientations, and reconstructions are inevitably limited by noise from various sources. At the same time, the field now benefits from an unprecedented wealth of high-quality density maps deposited in the public database^2^, providing a valuable resource that has yet to be fully exploited.

In practice, density maps obtained from experimental data may be degraded by noise, orientation bias, and reconstruction artifacts. Correcting these effects is an ill-posed inverse problem that requires more than the data alone. Bayesian inference offers a natural solution by combining prior knowledge about plausible structures with a likelihood term that captures dataset-specific statistics. Importantly, the iterative refinement process in single particle cryo-EM is highly non-convex, where noise and artifacts present in intermediate densities can propagate across iterations, bias pose assignments, and trap the optimization in suboptimal local minima, ultimately limiting the quality of the final reconstruction. Appropriately regularizing or denoising intermediate densities during refinement may reshape the effective optimization landscape, stabilize pose and density updates, and guide the reconstruction toward more physically plausible solutions. Recent efforts to improve reconstructions have largely taken the form of heuristic regularization^3,4^ or denoising strategies with pretrained models^5–8^, either within refinement or in post-processing. For example, Blush^6^ replaces RELION’s Gaussian prior^9^ with a pretrained denoiser learned from augmented training pairs of high-resolution maps. These approaches cannot incorporate dataset-specific statistics to guide or constrain the process. In addition, many depend on paired training data and may suffer from hallucinations or artifacts. Similar concerns extend to deep learning-based density modification tools^7,8,10^, which may produce hallucinated outputs and lose reliable low-frequency information.

From a Bayesian perspective, the ideal solution would be a data-driven prior that captures the statistical regularities of macromolecular structures directly from existing data. Generative models are naturally suited for this role, as they learn to represent the probability distribution of complex data. Diffusion models^11^ have recently shown remarkable success in high-fidelity generation across scientific and visual domains, while flow matching^12^ builds on the same intuition but provides a simpler and more efficient way to learn such transformations. In structural biology, generative models have been explored primarily in the atomic space for protein design^13–15^. Only a few recent studies have attempted to leverage these models to assist in the analysis and interpretation of experimental data^16–18^. However, since cryo-EM reconstruction operates directly in the density-map domain rather than atomic coordinates, such models cannot be readily applied, leaving many cryo-EM analyses without a suitable generative prior.

Here, we introduce cryoFM, a foundation model for cryo-EM densities that addresses these limitations by unifying prior knowledge and dataset-specific constraints within a modern generative framework. Trained in an unsupervised manner on thousands of high-quality cryo-EM maps using flow matching, cryoFM learns a powerful prior that captures the rich distribution of macromolecular densities. Combined with dataset-derived likelihood terms, it enables flow posterior sampling (FPS), an inference-only procedure where denoising and correction are explicitly constrained by dataset-specific statistics. This paradigm integrates seamlessly with existing expectation-maximization-style refinement loops while preserving Bayesian consistency, thereby enhancing interpretability. Furthermore, with only a small number of paired examples, cryoFM can be fine-tuned into conditional generative models for density modification. Unlike deterministic post-processing methods, these conditional models leverage the foundation model’s broad prior knowledge while retaining the controllability of posterior sampling, providing a flexible and reliable tool for practical applications.

### CryoFM learns generative priors of cryo-EM densities

CryoFM is a generative foundation model for cryo-EM densities that supports diverse downstream tasks. Using 3D U-Net^19,20^ as the model architecture (Extended Data Fig. 1), it is trained with flow matching on thousands of curated EMDB half maps (Fig. 1a), learning to transform samples from a simple Gaussian distribution into the complex data distribution of high-quality cryo-EM densities (Fig. 1b). Through this unsupervised pretraining, cryoFM captures a generalizable prior of macromolecular structures. This prior can be incorporated into a Bayesian framework by combining them with dataset-derived likelihood terms. Using flow posterior sampling (FPS), the model performs denoising and correction constrained by dataset-specific statistics, without additional training, and naturally extends to downstream tasks such as different modes of 3D refinement (Fig. 1c). With additional input-target training pairs, cryoFM can further be finetuned into conditional generative models that produce locally sharpened maps or maps simulated from atomic models (Fig. 1a), enabling task-specific density modification while retaining the controllability of posterior sampling (Fig. 1c).

**Figure 1.**
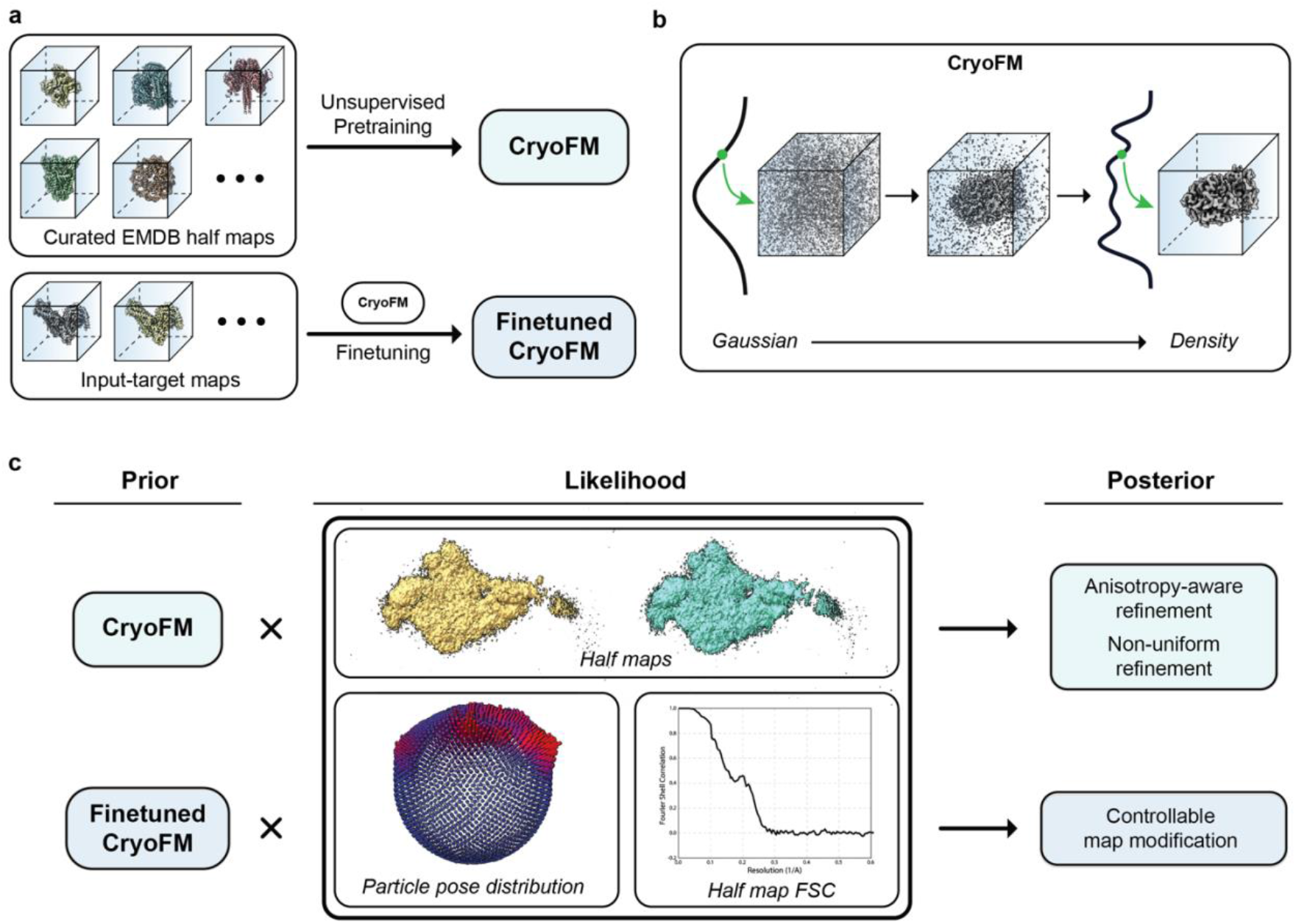
CryoFM is a generative foundation model that learns priors of cryo-EM densities and enables various downstream tasks, **(a)** CryoFM is pretrained in an unsupervised manner on curated EMDB half maps to capture general generative priors of high quality cryo-EM densities. The pretrained model can then be finetuned on specific input-output maps for task-specific adaptation, **(b)** CryoFM is a flow-based generative model that learns a continuous mapping from a simple Gaussian distribution to the complex data distribution of cryo-EM densities, **(c)** CryoFM or its finetuned variant can serve as a Bayesian prior in combination with likelihood terms derived from cryo-EM dataset-specific statistics. This integration enables downstream tasks such as anisotropy-aware or non-uniform refinement, or style-specific map modifications in a controllable manner.

### CryoFM enables Bayesian posterior sampling for cryo-EM inverse problems

FPS combines the generative prior from cryoFM with likelihood terms that describe how a high-quality density is transformed into the observed data (Fig. 2a). These likelihoods capture forward processes that map ideal densities to degraded observations, ranging from simple noise models to dataset-specific effects such as anisotropy from preferred orientations or other experimental distortions. Because they model degradation rather than restoration, such processes are often easier to specify and can be readily incorporated into FPS. In this formulation, the prior contributes structural information learned from high-quality maps, while the likelihood encodes dataset-specific statistics, yielding posterior densities that integrate both sources of information without additional training.

**Figure 2.**
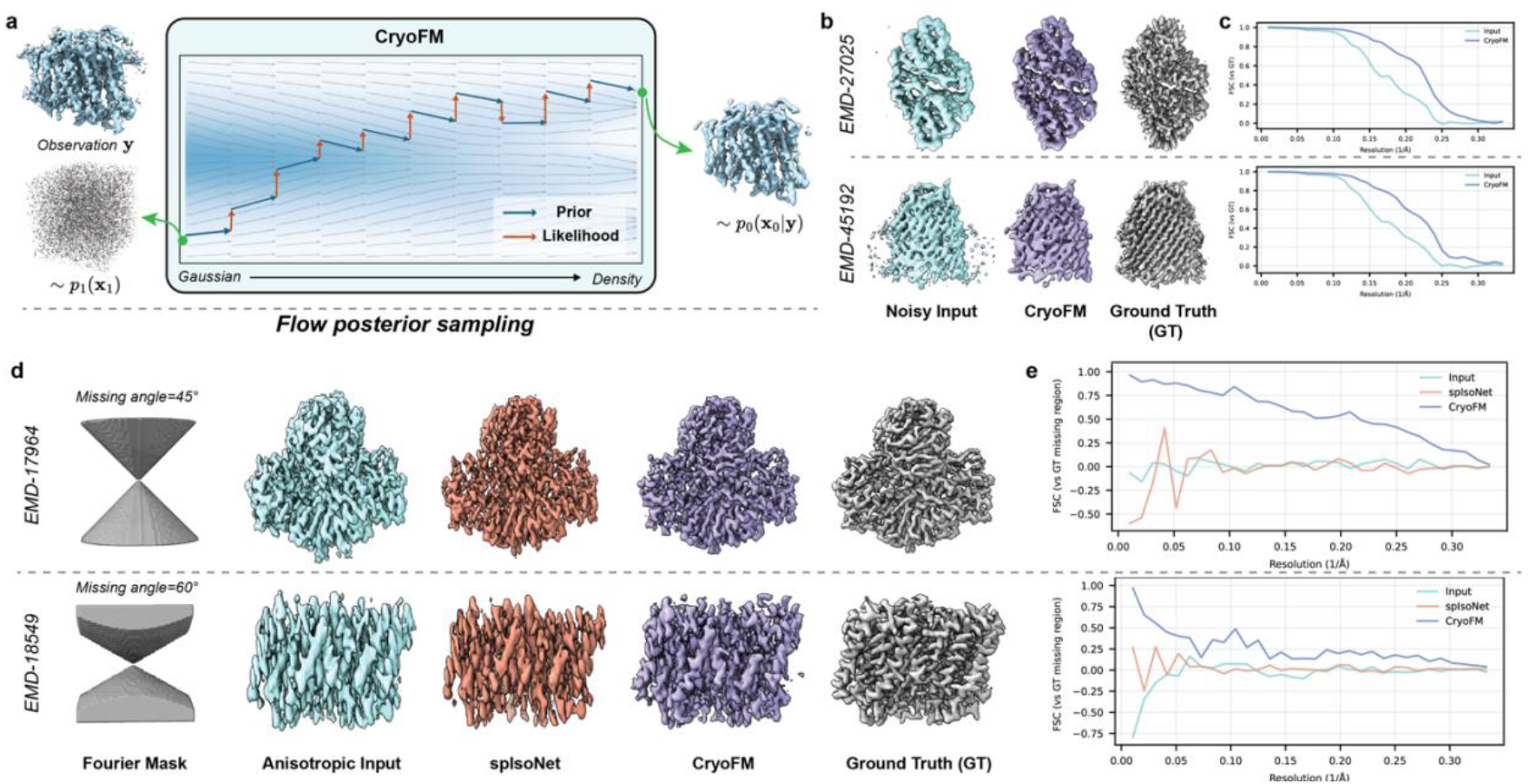
CryoFM enables data-constrained generative inference for cryo-EM densities, **(a)** Schematic of flow posterior sampling (FPS) with cryoFM. A generative prior learned by flow matching transforms samples from a Gaussian distribution into the density space of cryo-EM maps, while dataset-specific likelihood terms guide posterior sampling toward densities consistent with the observation. Blue arrows denote the learned prior flow and orange arrows indicate the likelihood-driven correction during inference, **(b)** CryoFM-based denoising under the synthetic noise setting. Input densities were synthetically corrupted and subsequently denoised using cryoFM. Both input and output densities were postprocessed with FSC-based filtering and B-factor sharpening using the same B factors for visualization, **(c)** FSC curves calculated between the noisy input and cryoFM denoised output against the ground truth density, **(d)** Anisotropy correction on densities with synthetically masked Fourier-space information. Directional Fourier-space masks was applied to the ground-truth densities to generate anisotropic inputs that mimic missing information from anisotropic sampling. SplsoNet and cryoFM were applied for restoration, with reconstructions shown alongside the ground truth, **(e)** FSC computed between the densities shown in **(d)** and the ground truth density, evaluated only within the synthetically masked Fourier-space region.

We first evaluated FPS in a synthetic denoising setting (Fig. 2b). Starting from deposited EMDB densities, we generated pairs of noisy half maps by adding frequency-dependent spectral noise, producing inputs with substantially reduced signal-to-noise ratio. These synthetically corrupted densities were then denoised using FPS with cryoFM as the generative prior. For visualization, both the noisy inputs and the cryoFM-denoised outputs were subjected to FSC-based frequency filtering and sharpened with identical B-factors. In this setting, cryoFM suppresses noise while preserving structural features (Fig. 2b). This improvement is supported by increased FSC agreement with the ground-truth densities across frequencies (Fig. 2c).

We next tested whether the same framework could address anisotropic information loss, a common challenge in cryo-EM arising from preferred particle orientations. To this end, we constructed a synthetic task in which anisotropic input densities were generated by masking a symmetric conical region in Fourier space, mimicking missing angular information (Fig. 2d). These anisotropic inputs were then restored using spIsoNet^21^, a self-supervised method for anisotropy correction, and cryoFM. Visually, cryoFM produces restored densities with reduced anisotropic artifacts (Fig. 2d and Extended Data Fig. 2). Quantitatively, we computed the FSC only within the masked region in the Fourier space between each restored density and the ground-truth density (Fig. 2e). Under this evaluation, cryoFM consistently achieves higher FSC values, indicating improved reconstruction of density features in regions absent from the input. Together, these experiments show that cryoFM can be reused as a generative prior within FPS to address different cryo-EM inverse problems without retraining.

### CryoFM enables improved refinement across different likelihood settings

Building on its ability to perform density denoising and correction, it is natural to integrate cryoFM directly into the refinement process. In the conventional expectation-maximization (EM) framework, refinement alternates between estimating particle alignments and reconstructing a 3D density^9^. Within this loop, cryoFM naturally serves as a prior distribution for the reconstructed density, while flow posterior sampling provides the restoration according to the likelihood function (Fig. 3a). This formulation preserves the statistical grounding of refinement, embedding a learned generative prior in a fully Bayesian manner.

**Figure 3.**
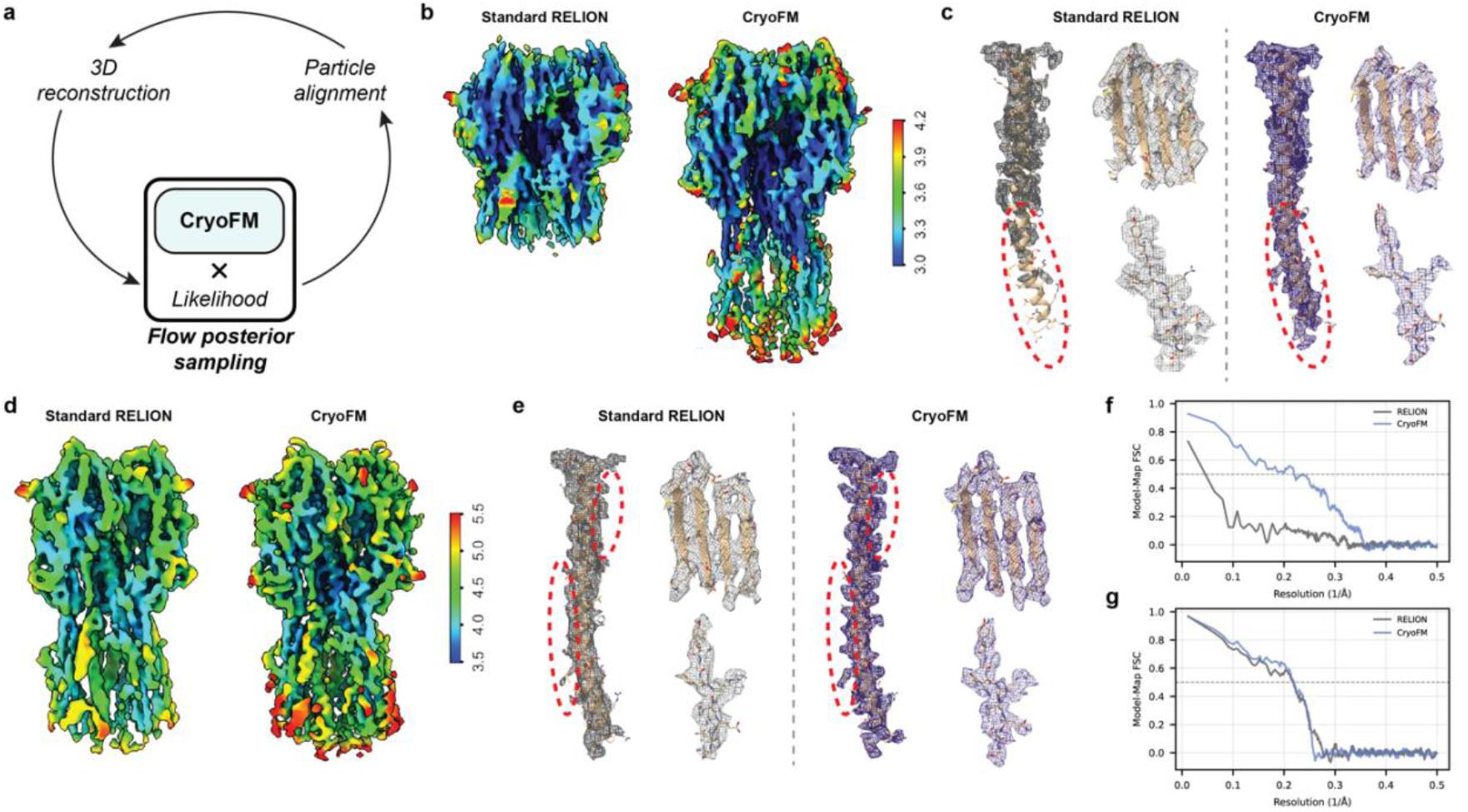
Anisotropy-aware refinement with cryoFM improves reconstruction quality relative to standard RELION, **(a)** Schematic of the refinement workflow integrating cryoFM into the standard RELION expectation-maximization (EM) iterations. CryoFM performs flow posterior sampling, guided here by a likelihood derived from the particle pose distribution, to mitigate artifacts induced by anisotropy and improve reconstruction, **(b-e)** Results on EMPIAR-10096 **(b-c)** and EMPIAR-10097 **(d-e).** Comparison of maps reconstructed using standard RELION (left) and cryoFM anisotropy-aware refinement (right), visualized with local resolution coloring (Å). CryoFM consistently yields visibly improved map quality and clearer structure features, **(f-g)** Model-map FSC curves of EMPIAR-10096 **(f)** and EMPIAR-10097 **(g)** CryoFM (blue) exhibits improved FSC across a broad resolution range relative to standard RELION (gray).

We first validated cryoFM on a benchmark dataset with well-behaved, highly symmetric signal characteristics (EMPIAR-11792), which contains seven single-particle cryo-EM datasets of the DNA protection during starvation protein (DPS) collected at different stage tilts^22^. We integrated cryoFM into the EM refinement loop in RELION by inserting an additional FPS step after each reconstruction. In this benchmark setting, the likelihood term was defined under a simple isotropic spectral noise model, and flow posterior sampling (FPS) was performed to restore the clean density. The reconstructed maps showed no visible artifacts, confirming the correctness and numerical stability of the approach (Extended Data Fig. 3).

Building on the anisotropy correction results above, we next incorporated cryoFM into the refinement process to assess whether the same framework improves reconstruction quality under anisotropic sampling. By combining particle pose distributions with half map statistics, the likelihood term was adapted to account for anisotropy in the data. To ensure reliable orientation alignment under anisotropic sampling, we adopted the low-resolution preservation strategy used in spIsoNet, retaining reference information up to 10 Å during each refinement iteration. Incorporating cryoFM under this anisotropy-aware formulation produced improved reconstructions compared with standard RELION for both influenza hemagglutinin (HA) trimer particle sets EMPIAR-10096 (Fig. 3b, c, f) and EMPIAR-10097 (Fig. 3d, e, g)^23^. For both datasets, we further compared cryoFM with Blush regularization^6^, cryoSPARC^24^, and spIsoNet^21^ (Extended Data Fig. 4a-d and Extended Data Fig. 5a-d), and calculated the FSC curves against a higher-quality HA trimer map (EMD-21954) from a different dataset (EMPIAR-10532)^25^ as well as model-to-map FSC using a docked atomic model (PDB: 6WXB) (Extended Data Fig. 4e and Extended Data Fig. 5e). In EMPIAR-10096, both cryoFM and spIsoNet produced clear structural features, and the FSC curves showed closer resemblance to EMD-21954 and the atomic model. However, anisotropy-related metrics derived from half maps, including FSO, Bingham curves^26^ and sphericity from 3DFSC^23^, favored Blush and spIsoNet. These differences were not fully supported by visual inspection or FSC curves against the higher-quality reference map and atomic model, suggesting that half map-based anisotropy metrics may not reliably reflect reconstruction quality in this setting. A similar pattern was observed for EMPIAR-10097, where Blush and spIsoNet achieved higher half map-based metrics. Although spIsoNet showed higher local resolution and stronger FSC agreement with external references (Extended Data Fig. 5), the resulting maps contained significant distortions, with loop regions being altered to β-sheet-like features (Extended Data Fig. 6). In contrast, cryoFM produced reconstructions with cleaner density and without apparent artifacts.

This refinement framework can be readily extended to other scenarios by adapting the likelihood to different degradation processes. For datasets exhibiting substantial spatial variation in signal-to-noise ratio, the likelihood can incorporate local noise statistics to reflect spatial heterogeneity, analogous in spirit to non-uniform refinement in cryoSPARC^3^. Using this formulation, we conducted systematic evaluations across nine particle sets^27–32^ and compared the results with standard RELION refinement, Blush regularization and cryoSPARC non-uniform refinement. Across these datasets, incorporating cryoFM consistently yielded reconstructions with improved map quality relative to standard refinement, and performance at least comparable to the best existing approaches (Extended Data Fig. 7-15). In the PfCRT dataset^27^ (EMPIAR-10330), cryoFM produced reconstructions with improved map quality and higher global and local resolutions relative to standard RELION (Fig. 4a-c and Extended Data Fig. 7). In the TFIIIC τA complex dataset^28^ (EMPIAR-11762), cryoFM yielded slightly improved resolution and clearer side-chain features (Fig. 4d-f and Extended Data Fig. 8). In the LRRC8A:D VRAC^29^ dataset (EMPIAR-12510), compared to standard RELION, cryoFM substantially improved the effective resolution with more clearly resolved structural features (Fig. 4g-i and Extended Data Fig. 11). In some particle sets, Blush regularization achieved slightly higher half map FSC values than cryoFM; however, these differences were not fully supported by visual map quality or model-to-map FSC curves, which do not depend on half map consistency (Extended Data Fig. 7, 10 and 11). This suggests that half map-based FSC metrics may not fully reflect reconstruction quality in these settings. Together, these results demonstrate that cryoFM provides a robust and flexible refinement strategy across diverse datasets. More broadly, the plug-and-play likelihood design enables straightforward extension to more complex forward models and refinement settings.

**Figure 4.**
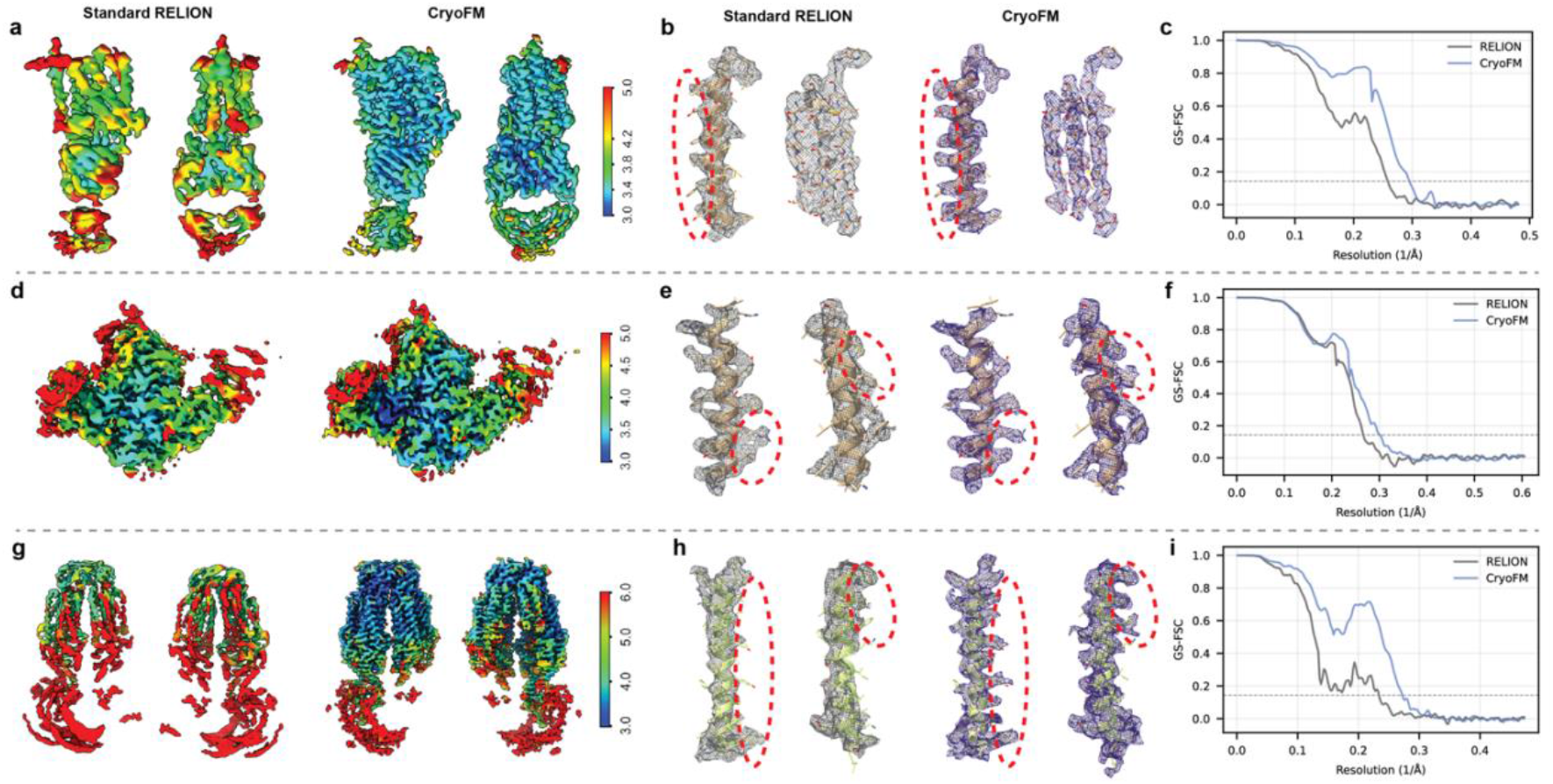
CryoFM non-uniform refinement improves reconstruction across benchmark datasets, **(a-i)** Comparison of density maps reconstructed using standard RELION (left) and cryoFM non-uniform refinement (right) for three datasets (EMPIAR-10330, EMPIAR-11762, and EMPIAR-12510, shown top to bottom), **(a, d, g)** Density maps visualized with local-resolution coloring (Å), showing overall improvements in map quality with cryoFM. **(b, e, h)** Zoom-in views of representative regions showing density overlapping with docked atomic models. CryoFM produces visibly sharper density at the detail level, including more clearly resolved side-chain features, demonstrating qualitative improvements beyond global resolution measures, **(c, f, i)** Half-map FSC curves showing that CryoFM achieves consistently higher FSC values across a broad resolution range compared to standard RELION

### Finetuned cryoFM offers controllable, different-style density post-processing

In addition to serving as an unconditional prior, cryoFM can be fine-tuned into conditional generative models for density post-processing (Fig. 5a). Using paired training data, we constructed two variants. The first was trained with half maps as input and LocScale-sharpened maps^33^ as targets, analogous to deep learning-based sharpening tools such as DeepEMhancer^8^. The second was trained with sharpened deposited maps as input and atomic model-simulated densities as targets, similar in spirit to EMReady^7^. Across both tasks, fine-tuned cryoFM outperformed the supervised baselines on most evaluation metrics when trained on the same datasets, particularly in map-model FSC (Fig. 5b and d, Extended Data Fig. 16a and Extended Data Fig. 17a). For real-space correlations (Extended Data Fig. 16b-e and 17b-e), cryoFM achieved higher values than DeepEMhancer, while EMReady obtained slightly higher values than cryoFM.

**Figure 5.**
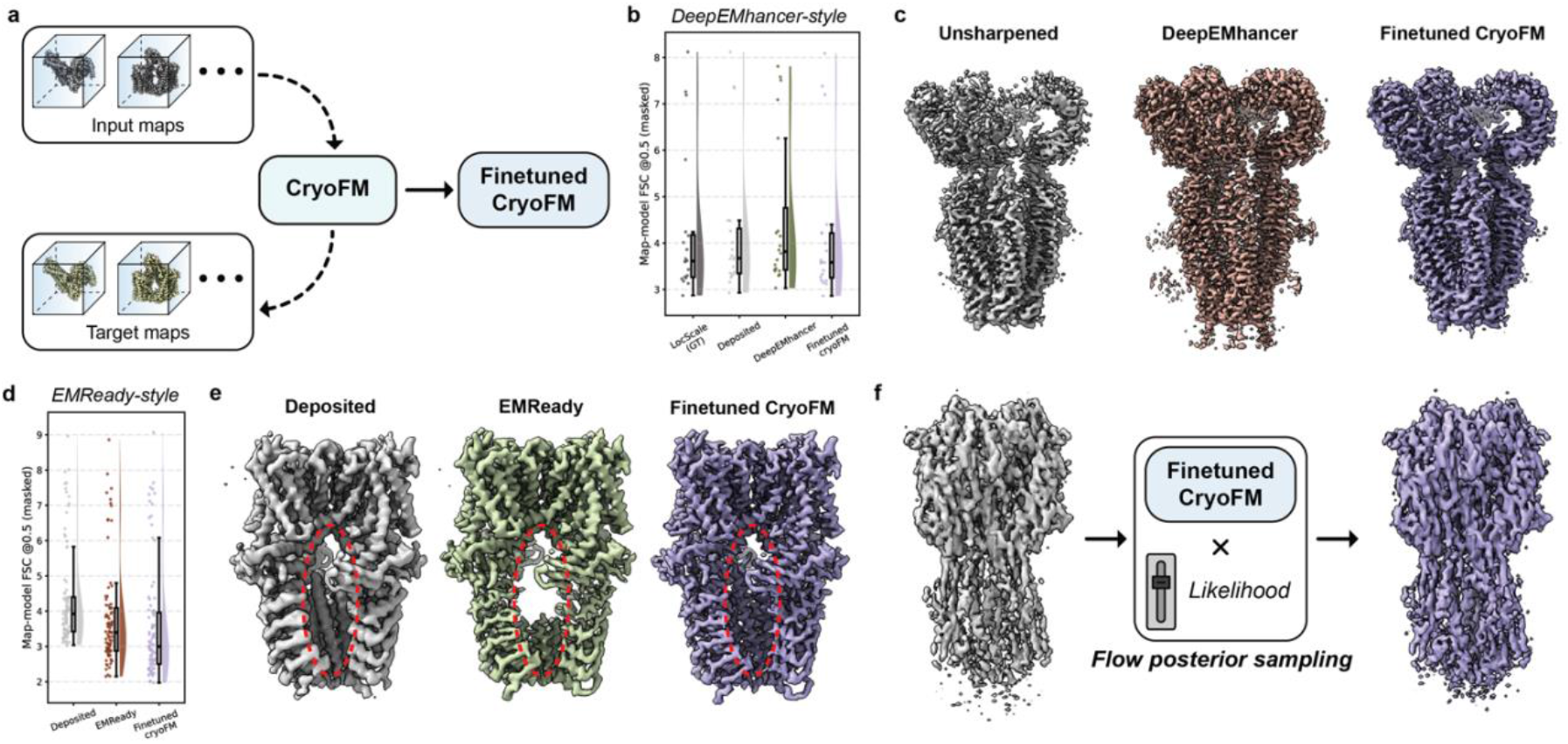
CryoFM enables controllable different style density post-processing through finetuning and likelihood-guided generation, **(a)** CryoFM is finetuned on paired input and target maps to adapt the pretrained foundation model for different style density post-processing, **(b)** Map-model FSC resolution at the 0.5 cutoff comparing the training target, input map, and maps post-processed by DeepEMhancer and fine-tuned cryoFM. **(c)** Representative density (EMD-6952) highlighting qualitative differences among the unsharpened map, DeepEMhancer output, and finetuned cryoFM. **(d)** Map-model FSC resolution at the 0.5 cutoff comparing the input map, EMReady output, and finetuned cryoFM post-processed maps, **(e)** Representative density (EMD-21128) illustrating qualitative differences among the input map, EMReady output, and finetuned cryoFM **(f)** Finetuned cryoFM used as a generative prior combined with an anisotropy-aware likelihood for flow posterior sampling, enabling controllable EMReady-style density modification on the EMPIAR-10096 reconstruction.

Beyond test-set metrics, fine-tuned cryoFM also produced density maps with better resolved features and fewer artifacts. In the DeepEMhancer-style setting, post-processed EMD-6952 by cryoFM exhibited reduced noise and improved density continuity compared with DeepEMhancer (Fig. 5c and Extended Data Fig. 16f). In the EMReady-style setting, post-processed density for EMD-21128 produced by cryoFM was substantially sharpened, with more structural features visible (Fig. 5e and Extended Data Fig. 17f), whereas the EMReady output showed pronounced artifacts, including missing α-helices (Fig. 5e). Across additional examples, cryoFM yielded densities with more clearly resolved secondary structures, side chains and DNA base pairs relative to the supervised baselines (Extended Data Fig. 17f). These results highlight the advantage of initializing from a foundation model trained on thousands of EMDB maps, which provides a stronger and more generalizable prior than training task-specific networks from scratch.

A key distinction of finetuned cryoFM is that it remains a generative model and can thus be used within the FPS framework. This property enables controllable density modification through adjustments in the likelihood formulation. By varying the weight of the likelihood term, or by altering its functional form, one can flexibly tune how much the output map adheres to dataset-specific information versus the learned prior (Fig. 5f). In practice, this controllability provides a major advantage: for example, we can control how strongly the final output adheres to the frequency-dependent constraints it encodes, allowing larger modifications in low-SNR regions while preserving high-SNR frequency components. For example, by reusing the likelihood describing local noise statistics, we can perform EMReady-style density modification in a more controllable manner (Fig. 5f). By further adjusting the weight of the likelihood term, we can tune how faithfully the output follows these constraints, and the resulting maps illustrate how varying this weight produces outputs with different degrees of adherence to the likelihood (Extended Data Fig. 16g-h and 17g-h). Together, these results establish fine-tuned cryoFM as a flexible and controllable framework for cryo-EM density post-processing.

## Conclusion and discussion

This work establishes cryoFM as the first generative foundation model operating directly in the density space of biomacromolecules. By uniting large-scale unsupervised pretraining on experimental cryo-EM maps with flow posterior sampling (FPS), cryoFM provides a unified Bayesian framework for cryo-EM reconstruction that is both generalizable across tasks and transparent in its statistical grounding. Across diverse inverse problems in cryo-EM, cryoFM consistently improves reconstruction quality by combining a learned generative prior with dataset-specific statistics through inference rather than deterministic prediction. Compared to existing deep learning-based models for density post-processing, cryoFM offers greater controllability and interpretability, as the relative influence of the prior and likelihood can be explicitly adjusted during posterior sampling. Together, these results demonstrate how probabilistic generative models can be directly integrated into refinement and post-processing workflows to improve the robustness, reliability, and physical plausibility of cryo-EM reconstructions. Compared to our earlier proof-of-concept study^34^, this work substantially expands both methodological development and practical application to real cryo-EM datasets.

This framework can be naturally extended to a wide range of cryo-EM and cryo-ET tasks. At the reconstruction level, it encompasses ab initio modeling, focused refinement, particle polishing, and the analysis of flexible heterogeneity. Importantly, these different stages correspond to distinct forward models and noise characteristics, and can therefore be accommodated within the same Bayesian formulation by adapting the likelihood term while keeping the generative prior unchanged. Beyond serving as an external restoration module, the framework also opens the possibility of deeper integration into the refinement process itself. In particular, learned generative priors could be incorporated directly into gradient-based optimization schemes, providing regularized updates that may improve optimization stability and accelerate convergence in highly non-convex refinement landscapes. Such integration would allow the prior to influence not only the quality of intermediate density estimates, but also the trajectory of the optimization process. At a broader level, density-based priors with sufficiently large receptive fields to capture global structural context rather than local patches may further benefit upstream tasks such as particle picking and segmentation in both single-particle cryo-EM and *in situ* cryo-ET. Finally, as generative models based on atomic representations continue to advance, density-based foundation models such as cryoFM provide a natural interface for bridging experimental density maps with atomic modeling, enabling joint inference under explicit physical and statistical constraints.

Beyond these specific applications, this study shows that generative priors can serve not only to design or generate realistic structures, but also to help process experimental data. The framework outlined here provides a basis for further advances. First, more powerful priors could capture structure at multiple resolutions and explicitly represent uncertainty. Second, advances in posterior sampling algorithms could accelerate inference and improve robustness to degradation operators with unknown parameters. Third, richer likelihood functions could better reflect the complexity of degradations inherent to experimental observations. We believe this work opens the door to a new paradigm in structural biology, where generative priors are not only tools for designing realistic models, but also powerful instruments for processing, refining, and interpreting experimental data.

## Methods

### Flow matching formulation in cryoFM

In cryoFM, flow matching provides a continuous-time generative formulation that transforms samples from a simple Gaussian distribution into the complex distribution of macromolecular densities through an ordinary differential equation (ODE)^34^. Here, we represent a 3D Coulomb-potential map as **x** ∈ ℝ^*D*×*D*×*D*^, where each element corresponds to the potential value at a voxel on a uniform grid. The collection of such maps defines the data space 𝒳: = *R*^*D*×*D*×*D*^. We denote by *p*_data_(**x**) the empirical distribution over macromolecular density maps. CryoFM seeks to generate new samples from this distribution by transforming a standard Gaussian distribution *p*_init_(·) into the data distribution *p*_data_(**x**).

Formally, flow matching models this transformation from the initial distribution *p*_init_ to the target distribution *p*_data_ as a deterministic evolution of a variable **x**_*t*_ along time *t* ∈ [0,1]. The dynamics follow an ODE:

Formally, flow matching models this transformation from the initial distribution *p*_init_ to the target distribution *p*_data_ as a deterministic evolution of a variable **x**_*t*_, where the time parameter is traversed in the reverse direction from t=1 to t=0. The dynamics follow an ODE:

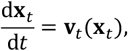

where **v**_***t***_(·): *R*^*D*×*D*×*D*^ → *R*^*D*×*D*×*D*^ is a time-dependent vector field, **x**_1_ ∼ *p*_init_ and **x**_0_ ∼ *p*_data_.

A simple and widely used choice is to define an explicit interpolation path between noise and data, for example the linear path:

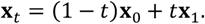

Along this path, the analytical velocity is:

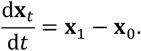

We train a neural network **v**_Θ_(*t*, **x**_*t*_) to approximate such velocity field from samples by minimizing the mean-squared error:

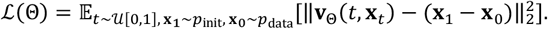

After training, integrating the learned ODE from *t* = 1 to *t* = 0 transforms Gaussian noise into a generated 3D density map that follows *p*_data_. This formulation avoids stochastic simulation as in diffusion-based models and serves as a data-driven prior for downstream tasks.

### Generative models as data-driven priors

In structural biology, experimental observations often provide incomplete or noisy information about the underlying molecular structure. Recovering a physically meaningful and biologically plausible reconstruction from such partial data requires incorporating prior knowledge as additional constraints. Traditional approaches encode prior knowledge through explicit regularization, such as enforcing smoothness^9^, suppressing high-frequency noise^3^, or restricting densities to be non-negative in cryo-EM reconstruction^24^. While effective in simple settings, these hand-crafted priors capture only limited, low-order patterns.

A generative model provides an alternative, data-driven route. Instead of hand-writing constraints, we learn the probability distribution *p*_data_(**x**) directly from the data. This data-driven prior model implicitly encodes what typical high-quality densities look like. Given a new experimental observation **y**, the prior *p*(**x**) can be combined with a physics-based likelihood *p*(**y**|**x**) under Bayes’ rule:

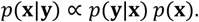

Here, *p*(**y**|**x**) encodes task-specific forward operators that describes the degradation process. The posterior *p*(**x**|**y**) thus balances data fidelity and biological plausibility, favoring densities that are both consistent with the measurements and typical under the learned distribution. In practice, we do not need to evaluate this posterior analytically; instead, we sample from it through flow posterior sampling (FPS), which couples the generative prior cryoFM with the task-specific likelihood in a unified inference procedure.

### Data preparation for pre-training

For model training, we constructed a dataset of cryo-EM density maps from the Electron Microscopy Data Bank (EMDB)^2^. To ensure high quality, we restricted the selection to entries resolved by single-particle cryo-EM with reported resolutions better than 3.0 Å and available half maps, such that resolution estimates were based on the gold-standard Fourier shell correlation (GS-FSC). We manually curated this set by excluding unusually large complexes, helical assemblies, and maps with severe artifacts or poorly resolved regions, as identified by visual inspection. Density maps with box sizes larger than 576 Å were also excluded. These exclusions were made to facilitate more effective model learning After curation, the dataset comprised 3,479 maps, of which 32 were set aside as a test set and excluded from training. The full list of EMDB entries used for training is provided in Supplementary Information Section C.

All selected maps were resampled to a unified voxel size of 1.5 Å using RELION^35^. To ensure a consistent numerical scale, density values were scaled by dividing them by their 99.999th percentile, bringing peak intensities to approximately 1. Subsequently, the maps were standardized based on global statistics derived from the protein regions of the training dataset (mean: 0.04; standard deviation: 0.09). This preprocessing reduced numerical variance and ensured stable model training. For training, each map was cropped into cubic sub-volumes of size 64^3^. These patches were then augmented using discrete 90-degree rotations and flips (via axis permutation). This approach produced 24 possible random orientations for each sample without requiring interpolation, thereby preserving the original voxel fidelity.

### Data preparation for DeepEMhancer-style post-processing

For the downstream task of DeepEMhancer-style post-processing, we constructed the training, validation, and test datasets strictly following the protocols and EMDB entry lists defined in the original DeepEMhancer paper. The training, validation and test sets contain 104, 21 and 20 maps respectively. We used input data consisted of half maps, paired with the corresponding target maps used in the original study. Notably, during data curation, we identified discrepancies where automatically generated masks were misaligned with the half maps; these cases were manually inspected and corrected using UCSF ChimeraX^36^ to ensure alignment accuracy. We first enforced a consistent Z-Y-X axis order in the MRC headers to ensure correct spatial orientation. Subsequently, all map pairs were resampled to 1.5 Å using RELION. Training preprocessing followed the pre-training strategy: maps were individually rescaled by their 99.999th percentile intensity, then standardized using the fixed global statistics (mean: 0.04, std: 0.09). We applied the same interpolation-free augmentation scheme (24 discrete orientations via axis permutation) during finetuning.

### Data preparation for EMReady-style post-processing

We utilized the publicly available EMReady dataset for this task. The training, validation and test sets contain 280, 70 and 90 maps respectively. All density maps were resampled to a pixel size of 1.5 Å using RELION. To ensure compatibility with Phenix validation tools and maintain accurate PDB-to-map alignment, we implemented a rigorous preprocessing protocol. First, we enforced a consistent Z-Y-X axis order and resampled all density maps to a unified pixel size of 1.5 Å. Next, to guarantee accurate coordinate handling in Phenix, we standardized the map origin definition by exclusively utilizing the *nstart* header attribute (ignoring the floating-point *origin* field). Since *nstart* requires integer grid indices, we used UCSF ChimeraX to resample the target (simulated) maps onto a grid where the physical origin aligned strictly with integer coordinates. Finally, the input maps (deposited) were resampled to match this geometry. For input maps, we applied the same pipeline: intensity scaling (division by the 99.999th percentile) followed by standardization (subtracting 0.04 and dividing by 0.09). For target maps, since they are simulated and inherently bounded within the [0, 1] range, they were directly standardized using the same statistics (mean: 0.04, std: 0.09). Data augmentation followed the same interpolation-free strategy as described previously.

### CryoFM model architecture

CryoFM is built on a 3D U-Net backbone that operates directly in the density space (Extended Data Fig. 1a). To enable conditional generation in later stages, the model is designed to accept a two-channel input tensor (with shape 2 × *D* × *D* × *D*). The first channel receives the noisy data patch 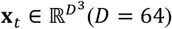, while the second channel accepts a conditioning tensor 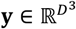. During pretraining, we set **y** = **0** (an all-zero tensor) to serve as a placeholder. The U-Net consists of four resolution levels, each containing two ResNet-style convolutional blocks with group normalization and SiLU activations^37^. Channel widths across the levels are set to (64, 128, 256, 512). Downsampling and upsampling between levels are implemented using convolutional layers. At the two deepest scales, self-attention layers^38^ are incorporated, with an attention head dimension of 8. Residual blocks at the middle scale further apply time-dependent modulation via positional time embeddings with a scale-shift mechanism^39^. Finally, a simple convolution layer maps the features back to the original data dimension, outputting the velocity field 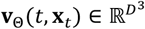.

### Finetuning cryoFM into conditional models

To enable controllable generation via classifier-free guidance (CFG)^40^, we extend the model input with a binary indicator channel 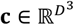 (Extended Data Fig. 1b). The network input is thus formed by concatenating the noisy data patch **x**_*t*_, the condition **y** and the indicator **c**. During fine-tuning, we apply a 10% dropout rate to the conditioning signal. For conditional samples, **y** contains the input density features and **c** is set to an all-one tensor (**1)**. For unconditional samples (dropout), both **y** and **c** are masked with all-zero tensors (**0**). This explicit indicator allows the network to dynamically switch between conditional and unconditional generation modes within the same training paradigm.

### Training details of cryoFM

CryoFM was pre-trained for 300,000 steps on 8 NVIDIA H100 GPUs with a global batch size of 96. The model was optimized using the AdamW optimizer^41^, with a peak learning rate of 1 × 10^−4^, betas (*β*_1_, *β*_2_) = (0.9, 0.98), and a weight decay of 0.01. The learning rate was linearly warmed up for the first 2,000 steps held constant thereafter. Training was performed in mixed precision (bfloat16) with gradient norm clipping set to 1.0. We maintained an exponential moving average (EMA) of the model parameters^11^ with a decay rate of 0.99, which was used for inference and evaluation.

For density modification applications such as DeepEMhancer and EMReady, we fine-tuned the base model separately for each task. Initialized from the pre-trained weights, each model was trained for an additional 300,000 steps using the identical hyperparameter setup (optimizer, learning rate schedule, and batch size) as in the pre-training phase.

### Flow posterior sampling (FPS)

Inverse problems in cryo-EM arise when one seeks to recover the clean density **x**_0_ from degraded observations **y** = 𝒜(**x**_0_), where 𝒜 is a forward operator modeling noise, filtering, or other experimental effects. Recent work has shown that pretrained diffusion models can be adapted to such problems via diffusion posterior sampling (DPS)^42^. DPS treats inverse problems by adding a likelihood score to the prior score of a pretrained diffusion model, and has been applied broadly without restrictive assumptions on the linearity of 𝒜. FPS mirrors this idea in the flow parameterization: instead of adjusting the prior score, we adjust the prior vector field by a likelihood gradient. Both views implement posterior-consistent sampling driven by a learned prior and a forward operator. In structural biology, this provides a natural route to process experimental data since most analysis tasks are inverse problems, in which the forward processes are typically easier to specify than explicit inverses, making FPS a practical and extensible tool.

In FPS, a pretrained flow model defines a vector field **v**_Θ_(*t*, **x**_*t*_) that transports a Gaussian distribution into the space of high-quality densities, modeling the prior distribution. To sample from the posterior, we adjust the flow field with the likelihood term:

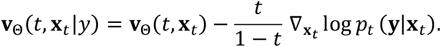

Substituting **x**_*t*_ in the likelihood term with Laplace approximation, where 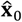 denotes the posterior mean estimated by the flow model, yields a practical update rule,

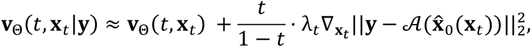

where the gradient enforces consistency with the observed data. In practice, we normalize this gradient for numerical stability and cap the weighting factor 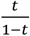 to prevent divergence. The resulting sampler integrates the generative prior with dataset-specific likelihoods, producing posterior-consistent reconstructions without additional training. More background, explanation and the algorithm of posterior sampling can be found in the Supplementary Information Section D.

### Task-specific forward operators

A central advantage of FPS is that it only requires modeling forward degradations that describe a high-quality density map to its observed counterpart, rather than an explicit inverse. This makes it straightforward to encode dataset-specific statistics. In this study, depending on the experimental condition, the forward operator used can represent spectral noise, anisotropic effects induced by preferred orientations, or local noise variations. Unlike restoration models, these operators only simulate how a clean density is transformed into an observation, making them easier to define and apply.

Spectral noise is the most widely used noise model in cryo-EM because it reflects the frequency-dependent noise characteristics introduced by the microscope optics. In this formulation, the forward operator is defined in the Fourier domain. Given the Fourier transform of a clean density map, ℱ(**x**) ∈ ℂ^*D*×*D*×*D*^, the observed volume is modeled as

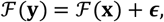

where *ϵ* is Gaussian noise with frequency-dependent variance. Specifically, for all Fourier coefficients on a spherical shell of radius *k*,

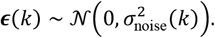

This captures the fact that higher frequencies generally exhibit greater noise variance than lower ones.

In practice, the distribution of particle orientations is often non-uniform, leading to anisotropy in the reconstructed density. This leads to frequency-dependent but also direction-dependent variations in the effective signal-to-noise ratio (SNR). Intuitively, regions of Fourier space that are under-sampled contain less reliable signal and thus higher noise variance, whereas directions with abundant sampling retain lower noise variance.

To capture this anisotropy, we assume that the radial noise variance estimated from FSC (as in the isotropic model) provides the baseline noise level at each spatial frequency shell. The directional distribution of particle sampling is then introduced as a modulation factor that redistributes this isotropic variance across orientations. This construction assumes that anisotropy primarily arises from uneven sampling, so that the radial average of the anisotropic variance matches the isotropic estimate.

Under the anisotropic noise setting, we model the observation in Fourier space as

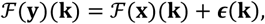

where ***ϵ***(**k**) is an additive complex Gaussian noise term whose variance depends on both frequency and direction,

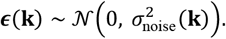

where ℱ(**x**) is the clean Fourier volume and **k** indexes a Fourier voxel with radial frequency *k* = ||**k**||.

The anisotropic noise variance is defined by modulating the isotropic radial baseline:

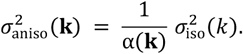

Here 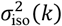 is the shell-wise noise variance estimated from half maps assuming isotropy, and *α*(**k**) is a direction-dependent modulation factor. Details of the derivation of *α*(**k**) can be found in Supplementary Information Section E.2.

In some cases, the noise in cryo-EM maps varies not only across spatial frequencies but also across different real-space regions. For example, membrane proteins often exhibit higher noise in the surrounding lipid layers compared to the protein core. To account for such spatial heterogeneity, we define a forward operator that captures local noise variance in real space while still preserving frequency sensitivity. Specifically, we construct a multiband representation based on the 3D Shannon wavelet transform^43^, which decomposes a density into a set of frequency bands while retaining its spatial locality. This formulation enables joint modeling of spatially and spectrally varying noise.

We model the observation in each wavelet band *b* as

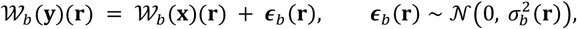

where 𝒲_*b*_(**y**)(**r**) and 𝒲_*b*_(**x**)(**r**) denote the observed and clean wavelet coefficients at spatial location **r**, and 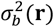 represents the local noise variance for band *b*. Each band captures fluctuations at a different frequency range, while the spatial dependence of 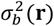 allows the model to adapt to regional variations in noise power.

### Details of the experiments

For synthetic denoising experiments, we started from averaged, unsharpened density maps in the EMDB test set. Frequency-dependent Gaussian noise with varying variance across spatial frequencies was added in Fourier space to generate two independently corrupted half maps. To prevent high-frequency noise from dominating real-space visualization, the corrupted half maps were subsequently weighted in Fourier space using frequency-dependent filtering. We then computed the half map FSC from the two synthetic half maps and used it to estimate the spectral noise variance. CryoFM was applied to each half map separately using flow posterior sampling (FPS), with the estimated noise statistics defining the likelihood term. For visualization and comparison, both the input (corrupted) and output (denoised) half maps were processed using FSC-based filtering and identical B-factor sharpening by `relion_postprocess`.

To evaluate restoration under anisotropic Fourier sampling, we constructed a symmetric missing cone mask in Fourier space (Fig. 2d). This mask was applied to averaged, unsharpened EMDB density maps by removing the corresponding Fourier coefficients, after which the masked volumes were transformed back to real space to simulate the effects of anisotropic sampling. CryoFM was then used to restore the masked densities using flow posterior sampling (FPS), with the known missing cone mask explicitly incorporated into the likelihood formulation.

All refinements with cryoFM were performed by integrating cryoFM into the RELION expectation-maximization (EM) algorithm using the `--external_reconstruct` flag. CryoFM-enabled flow posterior sampling (FPS) was applied from the first iteration and executed independently on the two half maps. The output half maps were subsequently filtered based on the calculated half-map FSC. To avoid introducing additional bias, FPS was not applied in the final iteration. For all particle sets, standard RELION was run using either the solvent mask provided in the corresponding EMPIAR dataset or a manually curated mask when no mask was available. The `--zero_mask` flag was omitted whenever its exclusion yielded improved reconstructions. Initial references were generated by low-pass filtering the deposited reconstruction to 40 Å. CryoFM, Blush regularization, and spIsoNet (when evaluated) were run with the same RELION parameters as their corresponding standard RELION baselines. In addition, Blush regularization was run with the spectral trailing enabled. CryoSPARC refinements were performed with default parameters and the dynamic masking enabled. For EMPIAR-10096, the `--keep_lowres` flag was used for the spIsoNet experiment.

For FPS, 200 sampling steps were used. Half maps were required to compute the likelihood term. Particle poses were included when anisotropy-aware likelihoods were used. An adaptive likelihood weight was employed to improve the robustness of generation across iterations. To reduce computational cost, each density was cropped to the bounding box defined by the solvent mask, and FPS was applied only to the densities within this region. Running FPS on six patches of size 64^3^ with a batch size of six takes approximately 44 seconds on a single V100 GPU.

For density modification experiments, the same test sets used by DeepEMhancer and EMReady were adopted to evaluate the finetuned cryoFM models. Classifier-free guidance (CFG) was used for conditional post-processing, with a CFG weight of 2.0 for DeepEMhancer-style map modification and 0.5 for EMReady-style map modification.

Global half map FSC curves, local resolutions, and the reported resolution cutoffs were computed in cryoSPARC with mask optimization enabled. Model-map FSC values and real-space correlation coefficients were obtained using Phenix^44^. The atomic models used for each dataset correspond to the structures originally deposited alongside the EMDB maps. FSO and Bingham statistics were computed using the implementations at https://github.com/Vilax/FSO/, and 3DFSC anisotropy analysis, including sphericity, was performed with default settings using the software at https://github.com/azazellochg/fsc3D.

## Extended Data Figures

**Extended Data Figure 1.**
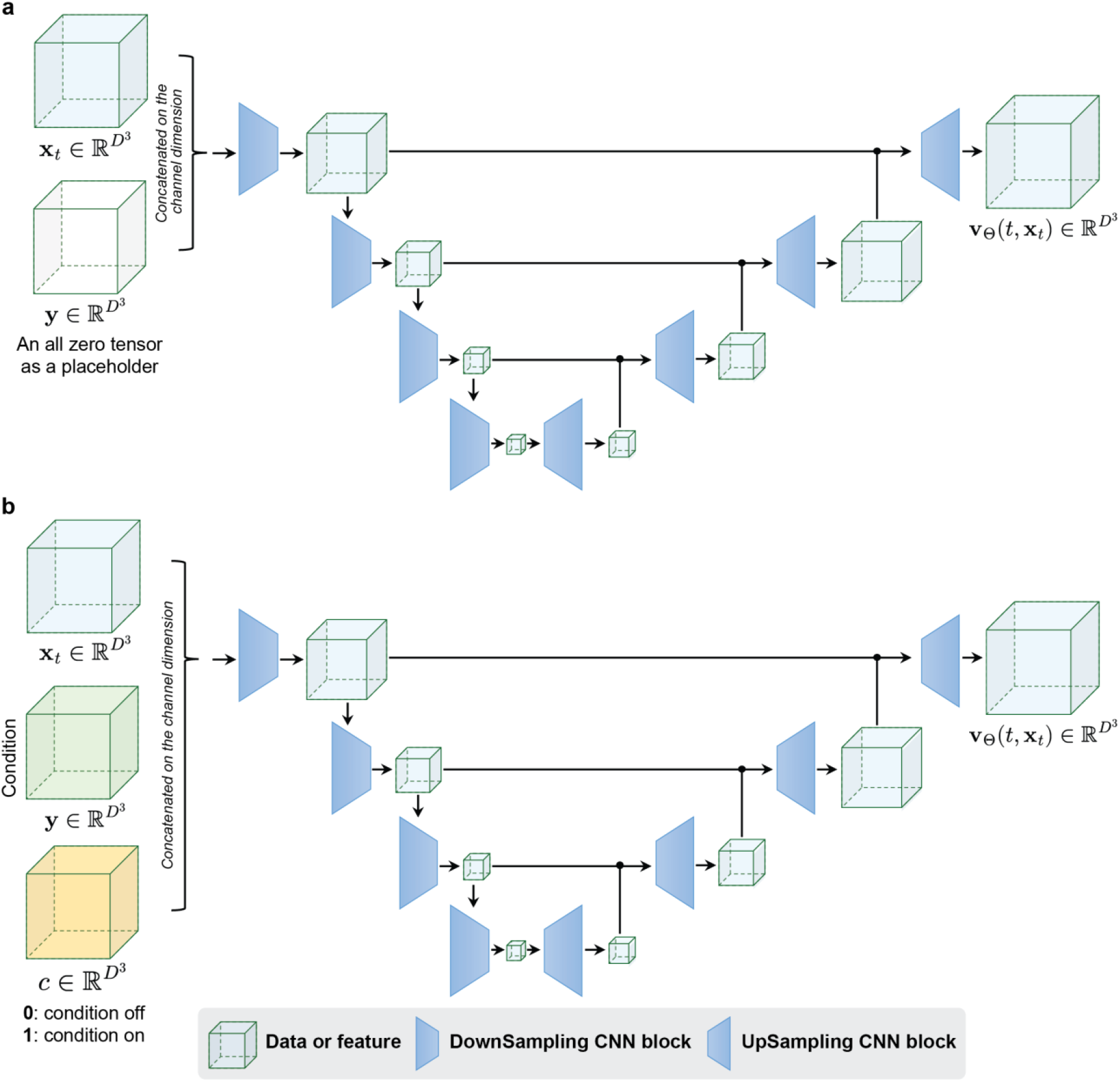
Network architectures used for pretraining and finetuning, **(a)** Pretraining architecture. The model takes a 2 × D × D × D tensor as input, where the second channel is an all-zero placeholder volume. The input is processed by a U-Net-style encoder-decoder with multi-scale downsampling and upsampling CNN blocks to estimate the vector field, **(b)** Finetuning architecture. An additional volume *c* is concatenated with the original inputs along the channel dimension, forming a 3 × D × D × D input tensor. This channel supports classifier-free finetuning, taking all zeros when no condition is used and all ones when a condition is present. The second channel corresponds to the conditioning density map. The same backbone architecture is used to predict the vector field.

**Extended Data Figure 2.**
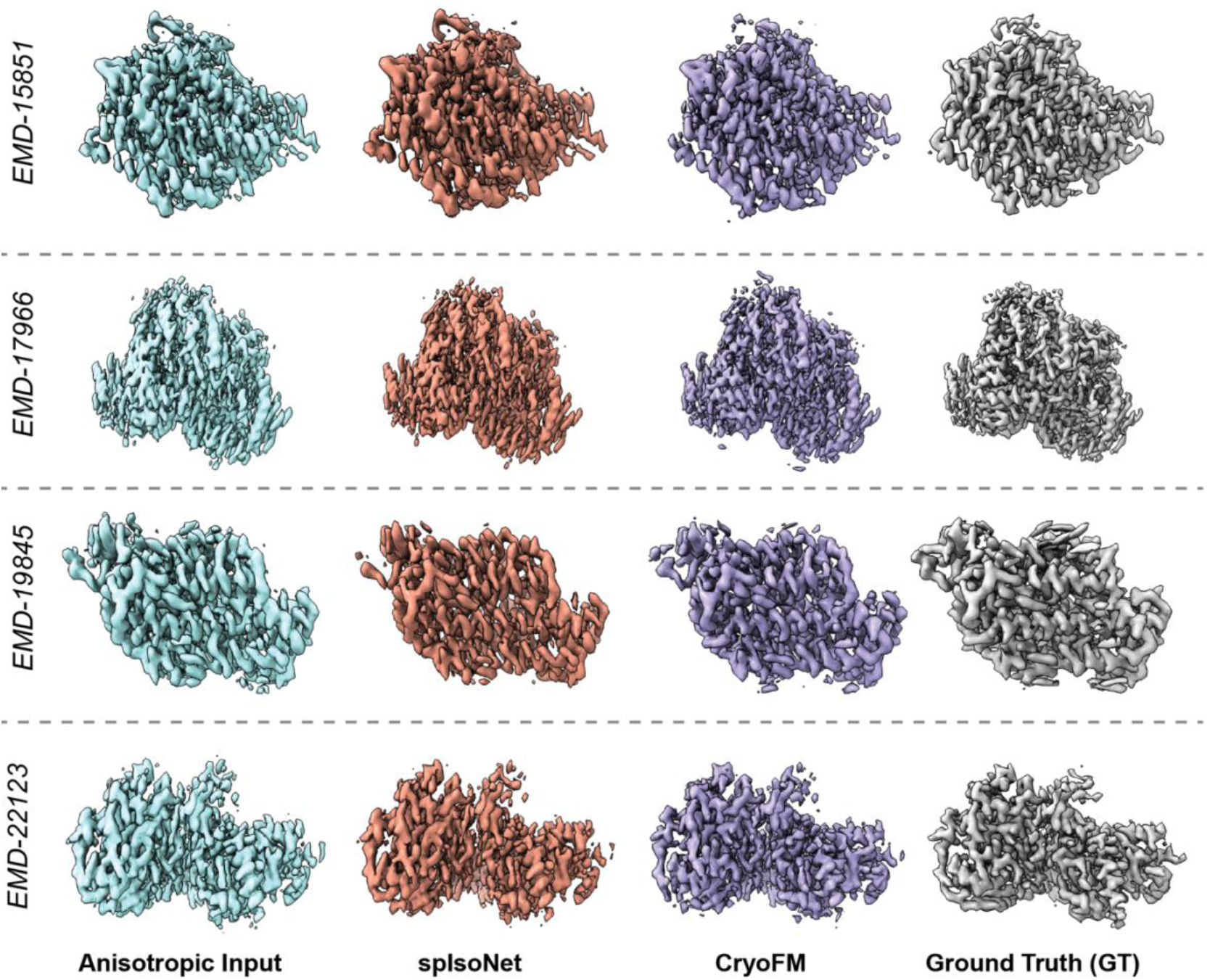
Anisotropy correction on EMDB densities after synthetic Fourier-space masking (missing cone angle is 45 degree). Directional Fourier-space masking was applied to ground-truth densities to generate anisotropic inputs. AR-Decon, spIsoNet and CryoFM were used for restoration, with results shown alongside the ground truth.

**Extended Data Figure 3.**
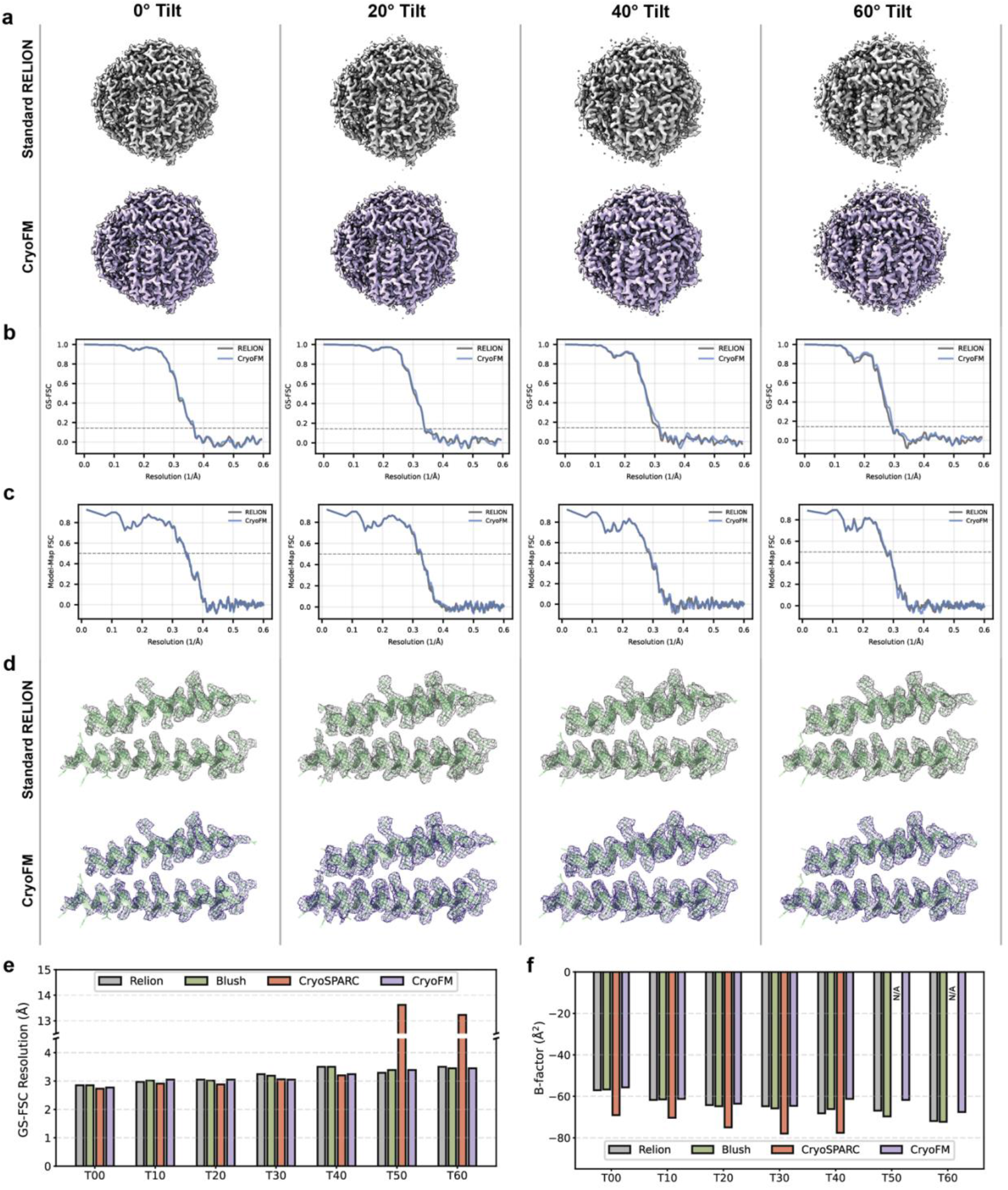
Performance of refinement methods across tilt conditions for EMPIAR-11**792. (a)** Density maps reconstructed using standard RELION (top row) and cryoFM (bottom row) under different tilt conditions (0°, 20°, 40°, 60°). (b) Half-map FSC curves and **(c)** model-map FSC curves of RELION and cryoFM across the four tilt angles, **(d)** Representative zoom-in density regions displayed with docked atomic models, **(e)** Summary of FSC resolution and **(f)** B-factor estimates calculated from RELION, Blush, cryoSPARC, and cryoFM refinement at each tilt setting (0°-60°).

**Extended Data Figure 4.**
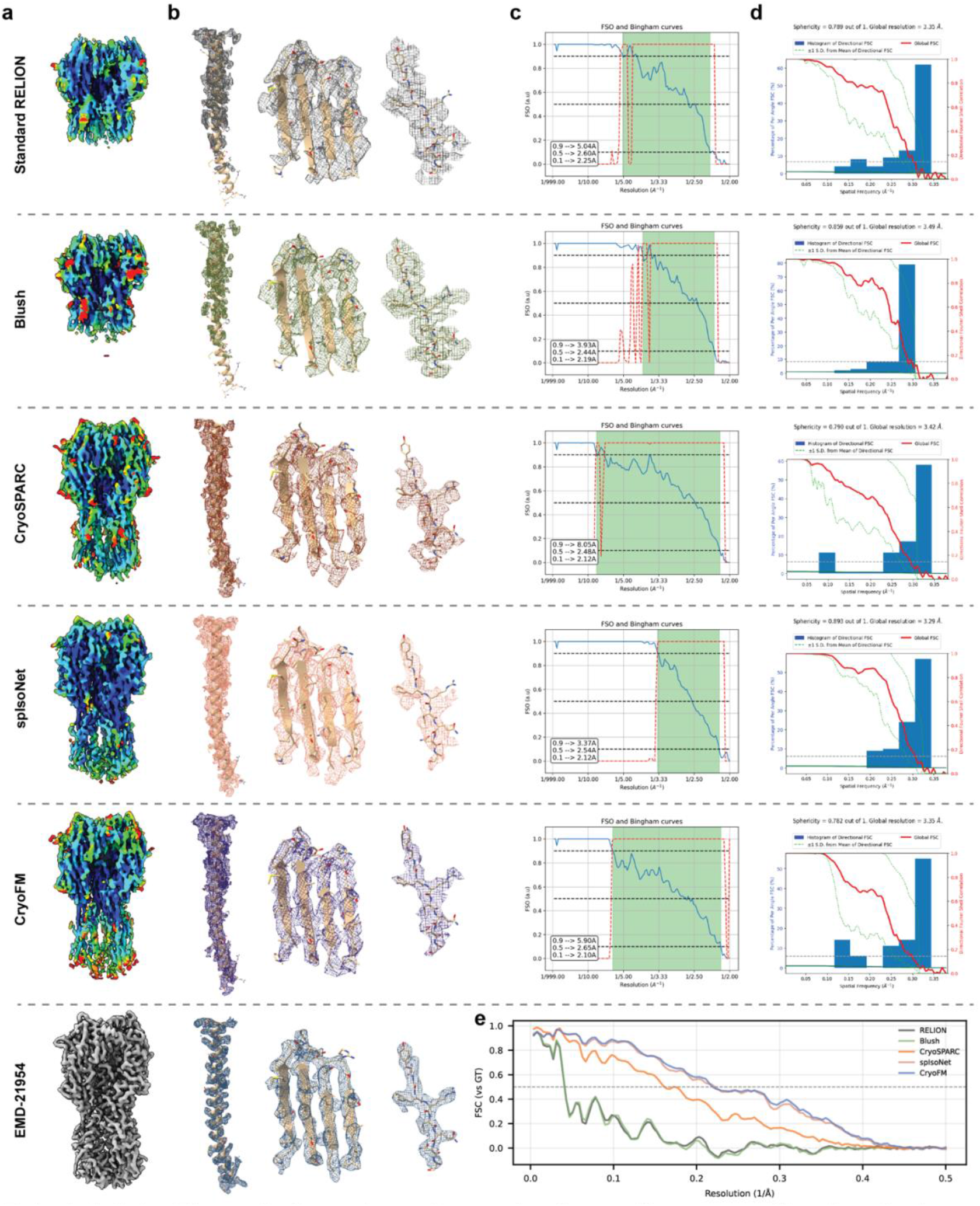
Comparison of reconstruction quality across different methods on EMPIAR-10096. Rows correspond to to standard RELION, Blush, cryoSPARC, splsoNet, cryoFM, and a high-quality reference map (EMD-21954) reconstructed from a different dataset used as ground truth (GT). Columns represent different visualization and quantitative assessment modalities, **(a)** Density maps colored by local resolution (Å), **(b)** Representative zoom-in density regions overlapping with docked atomic models, **(c)** FSO and Bingham curves, illustrating high-resolution feature recovery under each method, **(d)** 3DFSC plots and sphericity indicating anisotropy and directional resolution distribution, **(e)** FSC curves calculated between each method and the GT reference.

**Extended Data Figure 5.**
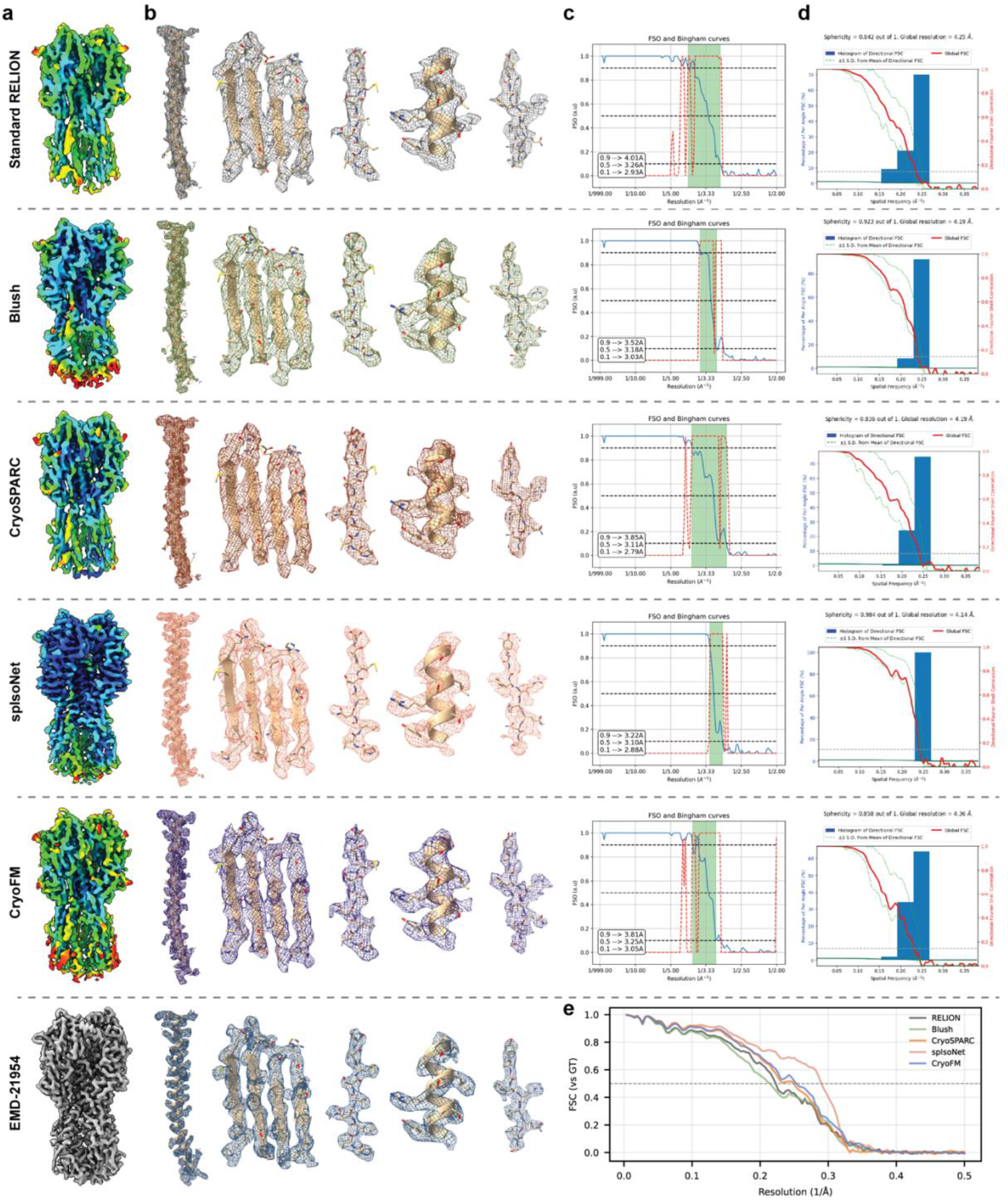
Comparison of reconstruction quality across different methods on EMPIAR-10097. Rows correspond to to standard RELION, Blush, cryoSPARC, spIsoNet, cryoFM, and a high-quality reference map (EMD-21954) reconstructed from a different dataset used as ground truth (GT). Columns represent different visualization and quantitative assessment modalities, **(a)** Density maps colored by local resolution (Å), **(b)** Representative zoom-in density regions overlapping with docked atomic models, **(c)** FSO and Bingham curves, illustrating high-resolution feature recovery under each method, **(d)** 3DFSC plots and sphericity indicating anisotropy and directional resolution distribution, **(e)** FSC curves calculated between each method and the GT reference.

**Extended Data Figure 6.**
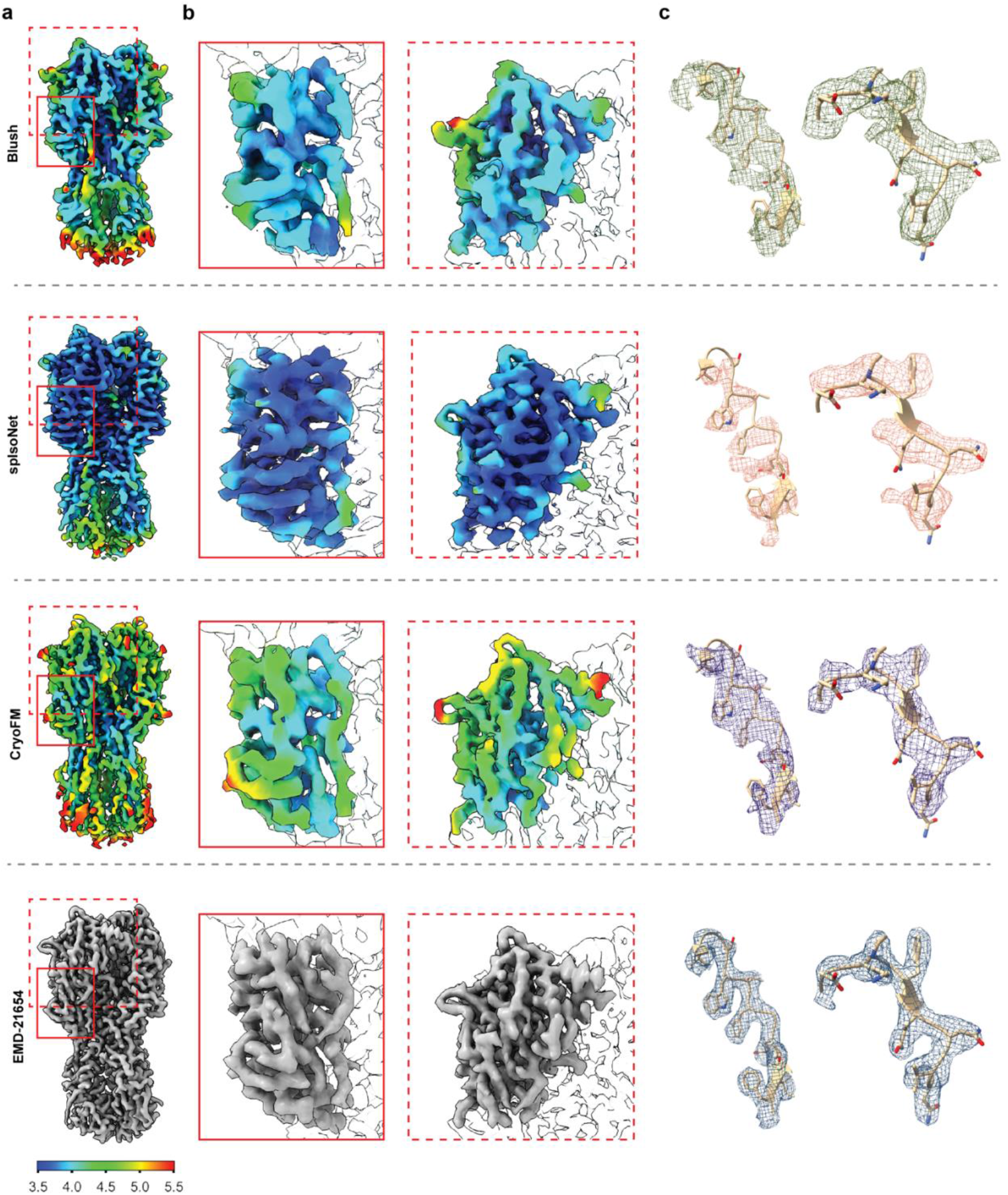
Comparison of structural artifacts across refinement methods. Visualization of density maps reconstructed using Blush, spIsoNet, and cryoFM, and a high-quality reference map (EMD-21654) used as ground truth (GT), **(a)** Full maps (colored by local resolution) with regions selected for detailed inspection highlighted by red boxes, **(b)** Zoom-in density views showing local map morphology, highlighting the structural artifacts in spIsoNet reconstructions, **(c)** Additional density regions displayed with docked atomic models to compare detail density features.

**Extended Data Figure 7.**
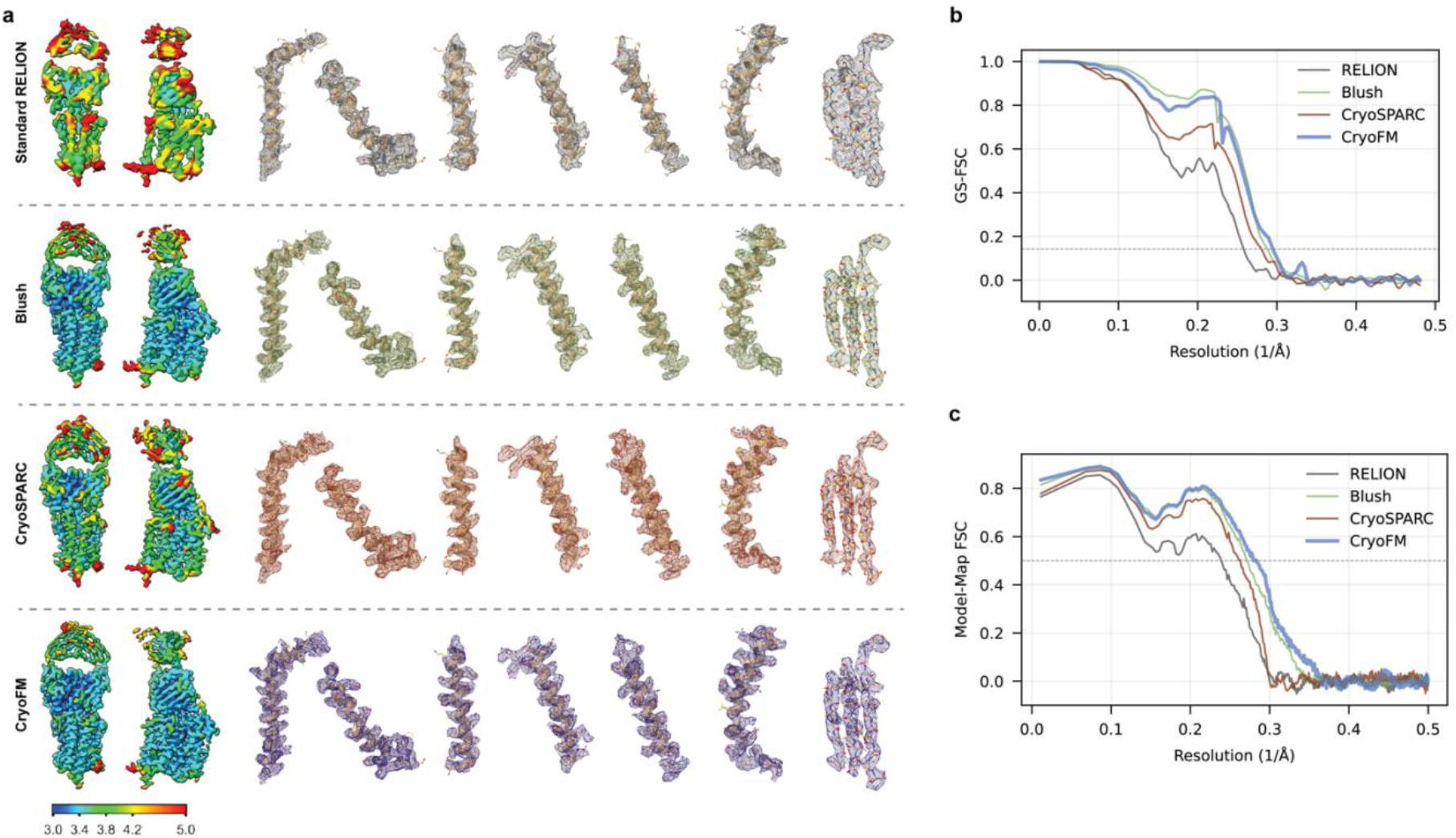
Comparison of refinement methods on EMPIAR-10330. **(a)** Density maps reconstructed using standard RELION, Blush, cryoSPARC, and cryoFM, shown with local resolution coloring (left) and representative zoom-in regions displayed with docked atomic models (right), **(b)** Half-map FSC curves and **(c)** model-map FSC curves of the four methods.

**Extended Data Figure 8.**
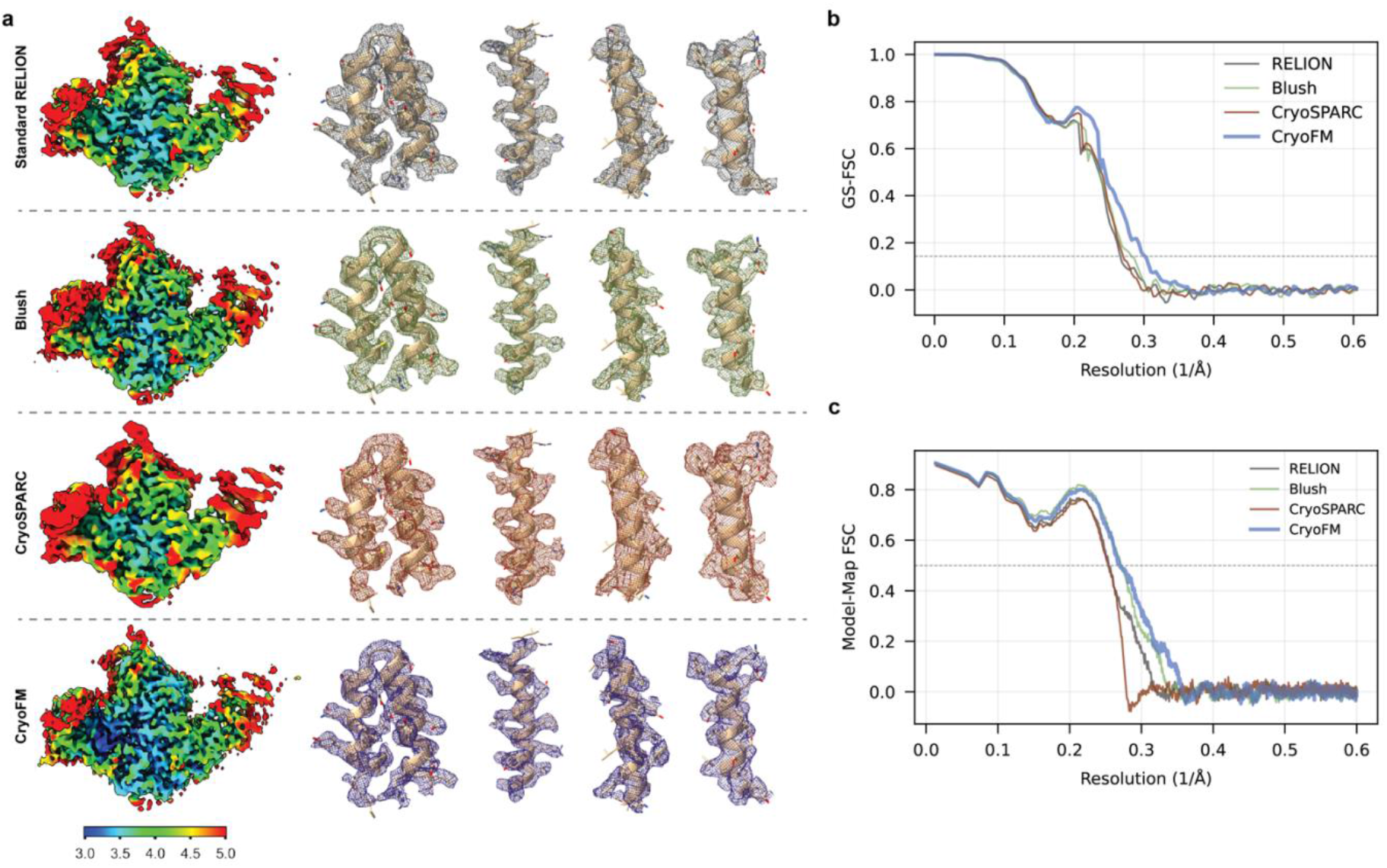
Comparison of refinement methods on EMPIAR-11762 (TauA). **(a)** Density maps reconstructed using standard RELION, Blush, cryoSPARC, and cryoFM, shown with local resolution coloring (left) and representative zoom-in regions displayed with docked atomic models (right), **(b)** Half-map FSC curves and **(c)** model-map FSC curves of the four methods.

**Extended Data Figure 9.**
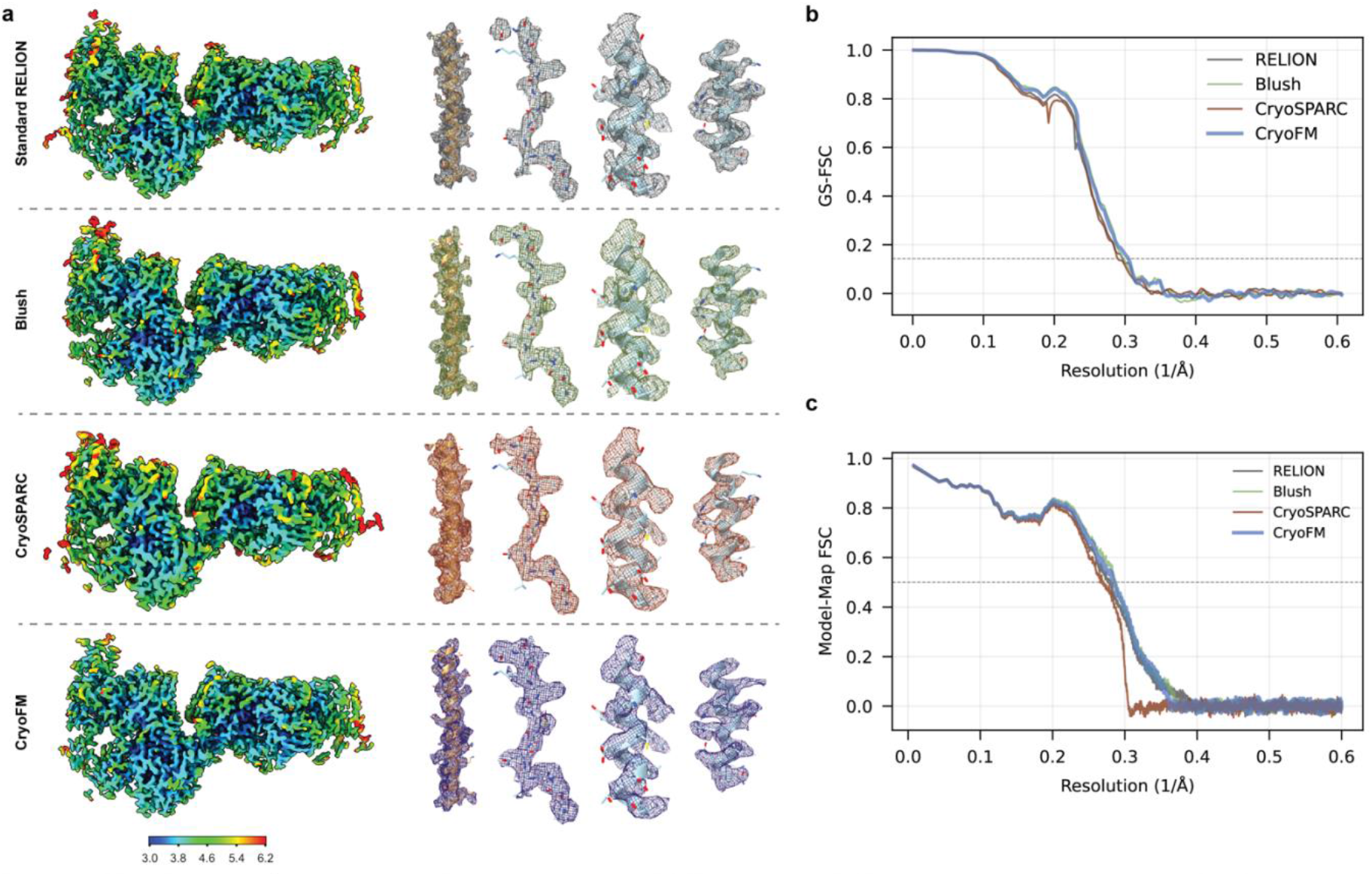
Comparison of refinement methods on EMPIAR-11762 (TauB). **(a)** Density maps reconstructed using standard RELION, Blush, cryoSPARC, and cryoFM, shown with local resolution coloring (left) and representative zoom-in regions displayed with docked atomic models (right), **(b)** Half-map FSC curves and **(c)** model-map FSC curves of the four methods.

**Extended Data Figure 10.**
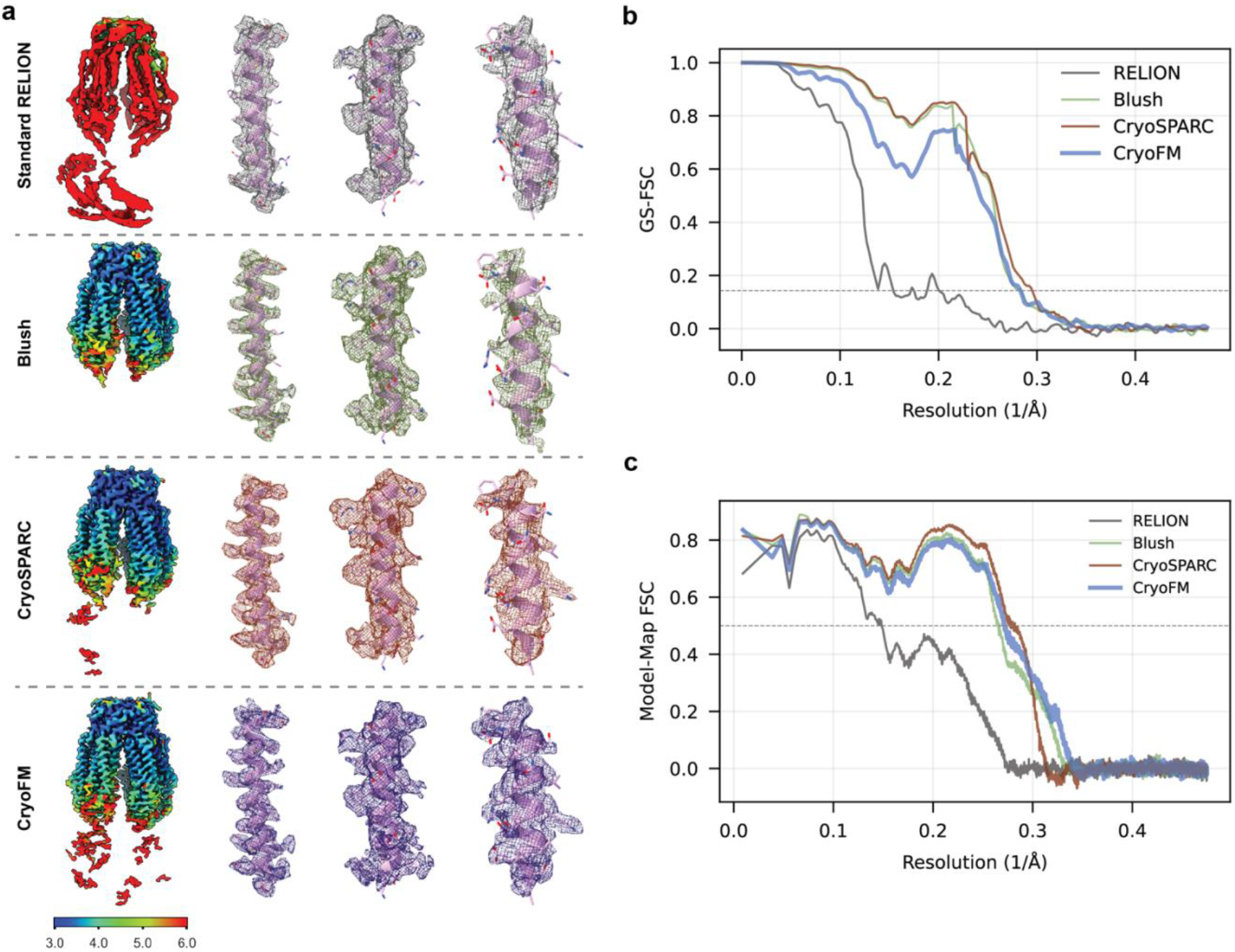
Comparison of refinement methods on EMPIAR-12510 (conformation 1). **(a)** Density maps reconstructed using standard RELION, Blush, cryoSPARC, and cryoFM, shown with local resolution coloring (left) and representative zoom-in regions displayed with docked atomic models (right), **(b)** Half-map FSC curves and **(c)** model-map FSC curves of the four methods.

**Extended Data Figure 11.**
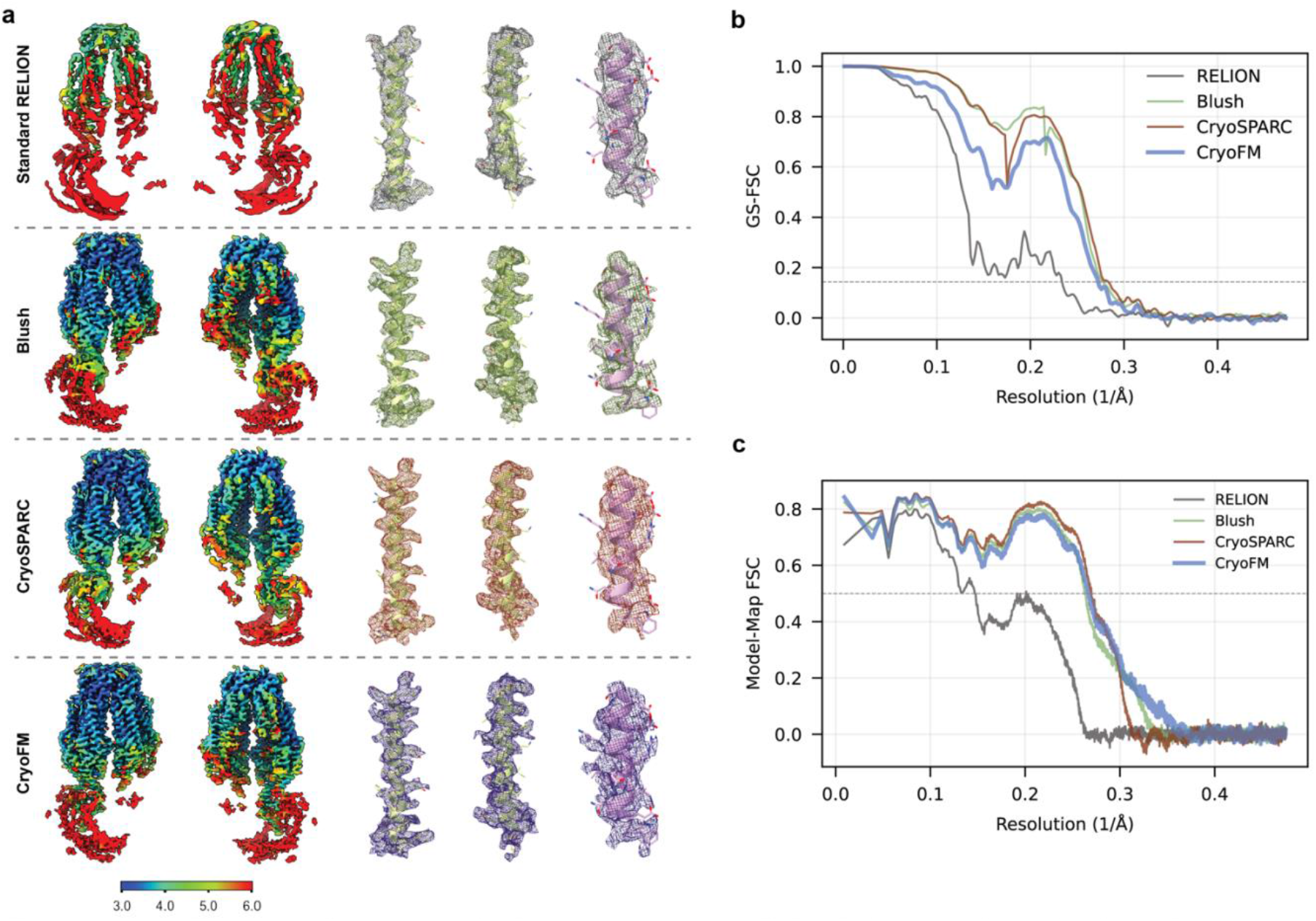
Comparison of refinement methods on EMPIAR-12510 (conformation 2). **(a)** Density maps reconstructed using standard RELION, Blush, cryoSPARC, and cryoFM, shown with local resolution coloring (left) and representative zoom-in regions displayed with docked atomic models (right), **(b)** Half-map FSC curves and **(c)** model-map FSC curves of the four methods.

**Extended Data Figure 12.**
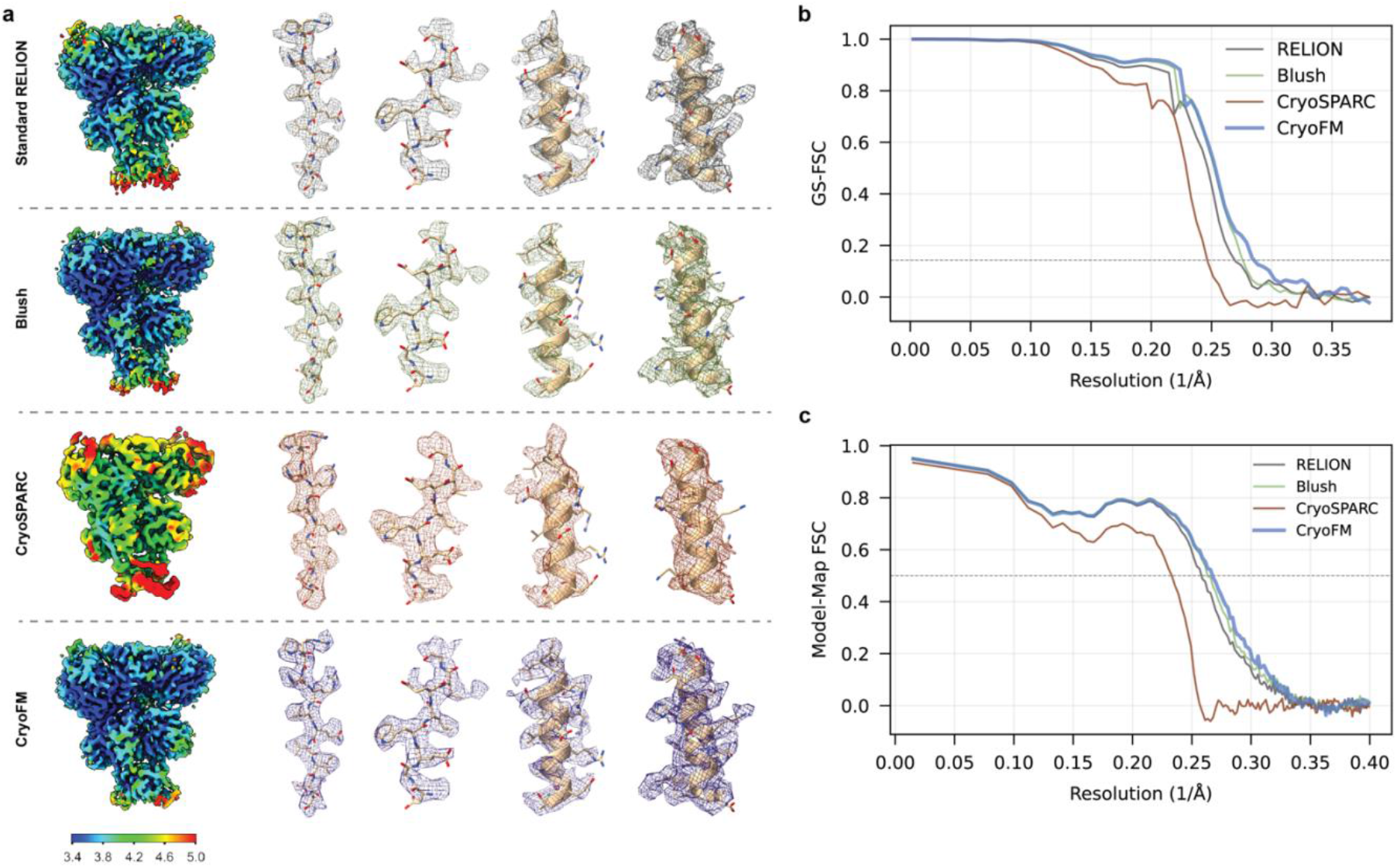
Comparison of refinement methods on EMPIAR-11247 (EMD-27645). **(a)** Density maps reconstructed using standard RELION, Blush, cryoSPARC, and cryoFM, shown with local resolution coloring (left) and representative zoom-in regions displayed with docked atomic models (right), **(b)** Half-map FSC curves and **(c)** model-map FSC curves of the four methods.

**Extended Data Figure 13.**
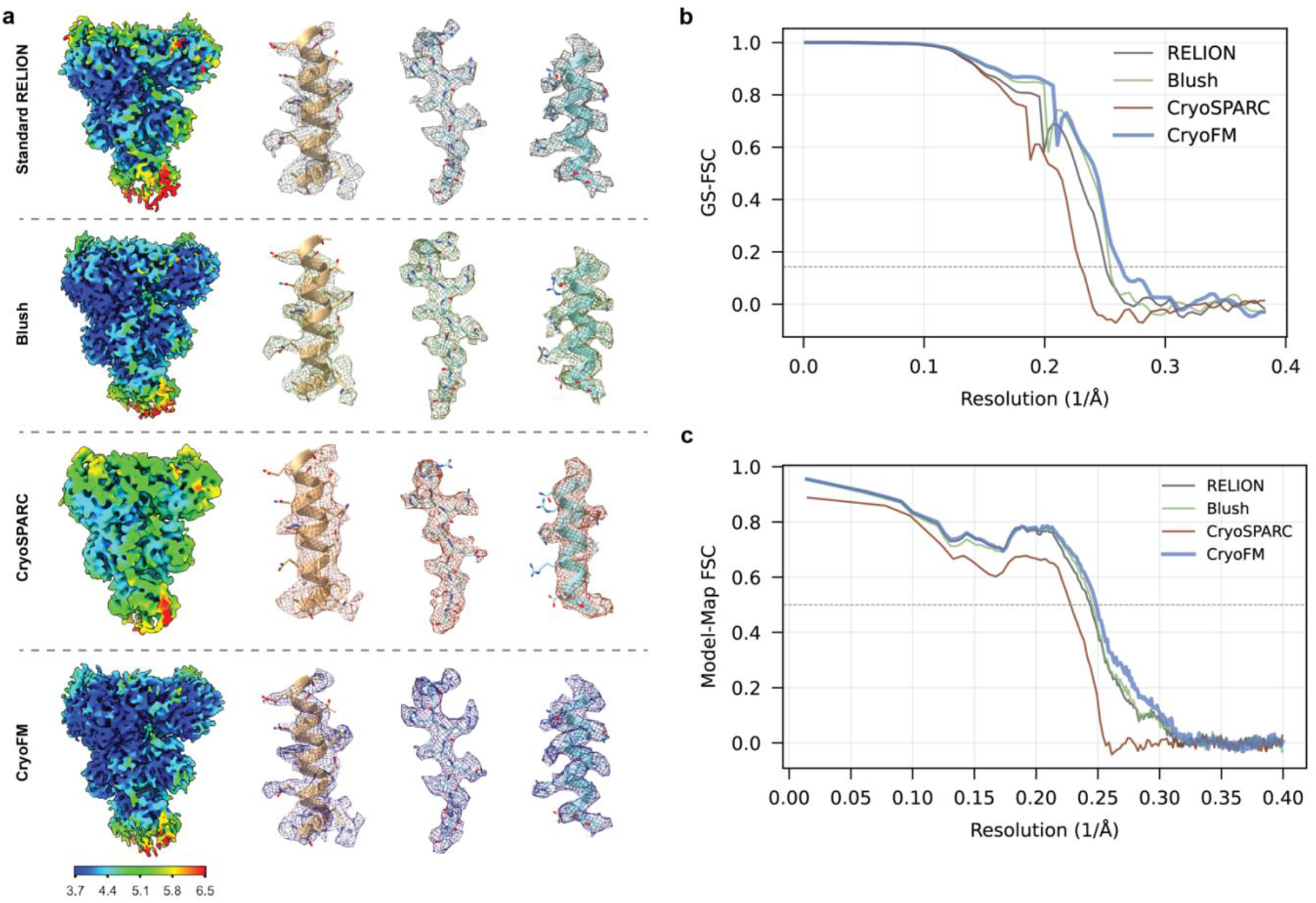
Comparison of refinement methods on EMPIAR-11247 (EMD-27646). **(a)** Density maps reconstructed using standard RELION, Blush, cryoSPARC, and cryoFM, shown with local resolution coloring (left) and representative zoom-in regions displayed with docked atomic models (right), **(b)** Half-map FSC curves and **(c)** model-map FSC curves of the four methods.

**Extended Data Figure 14.**
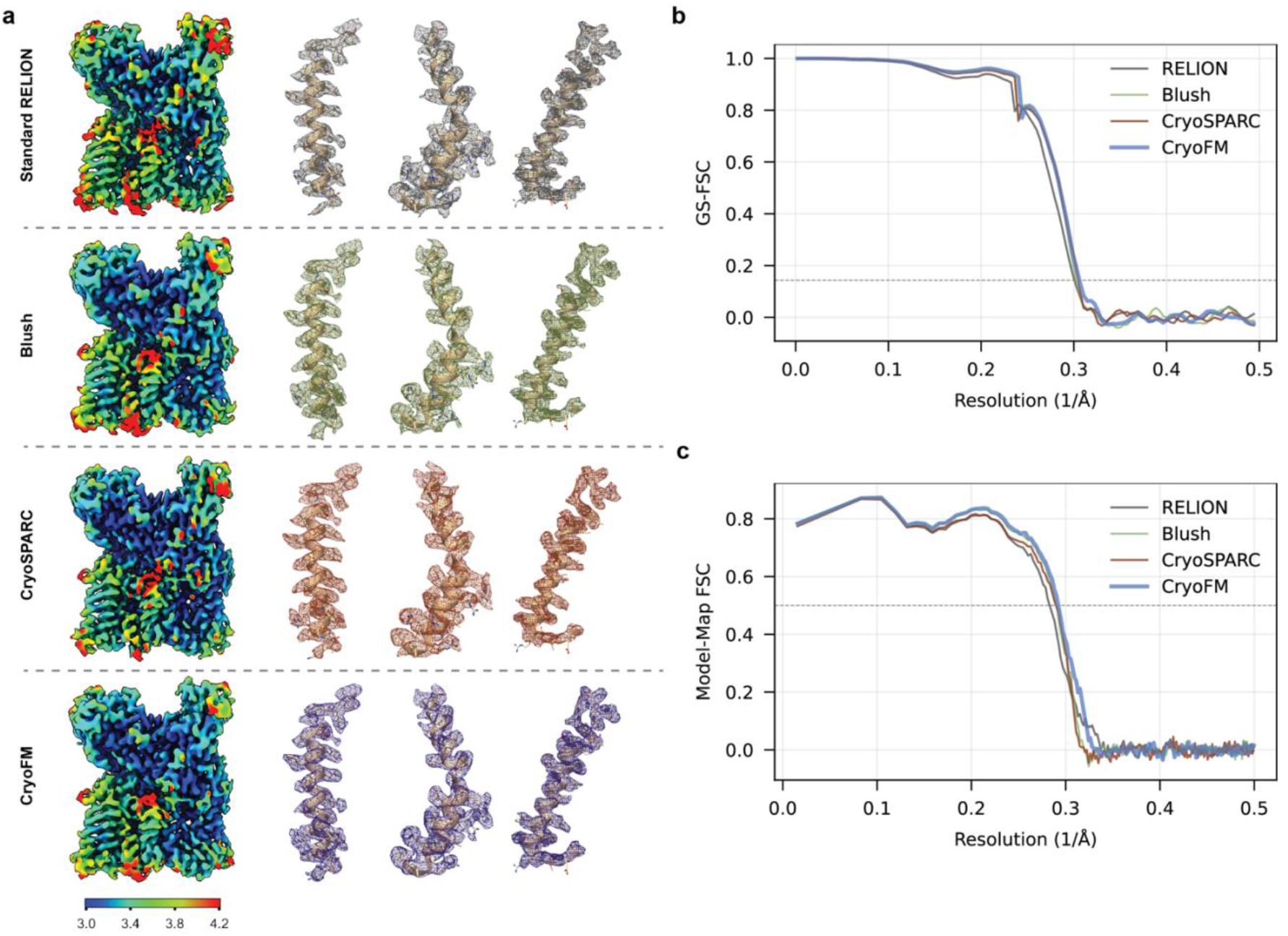
Comparison of refinement methods on EMPIAR-10420. **(a)** Density maps reconstructed using standard RELION, Blush, cryoSPARC, and cryoFM, shown with local resolution coloring (left) and representative zoom-in regions displayed with docked atomic models (right), **(b)** Half-map FSC curves and **(c)** model-map FSC curves of the four methods.

**Extended Data Figure 15.**
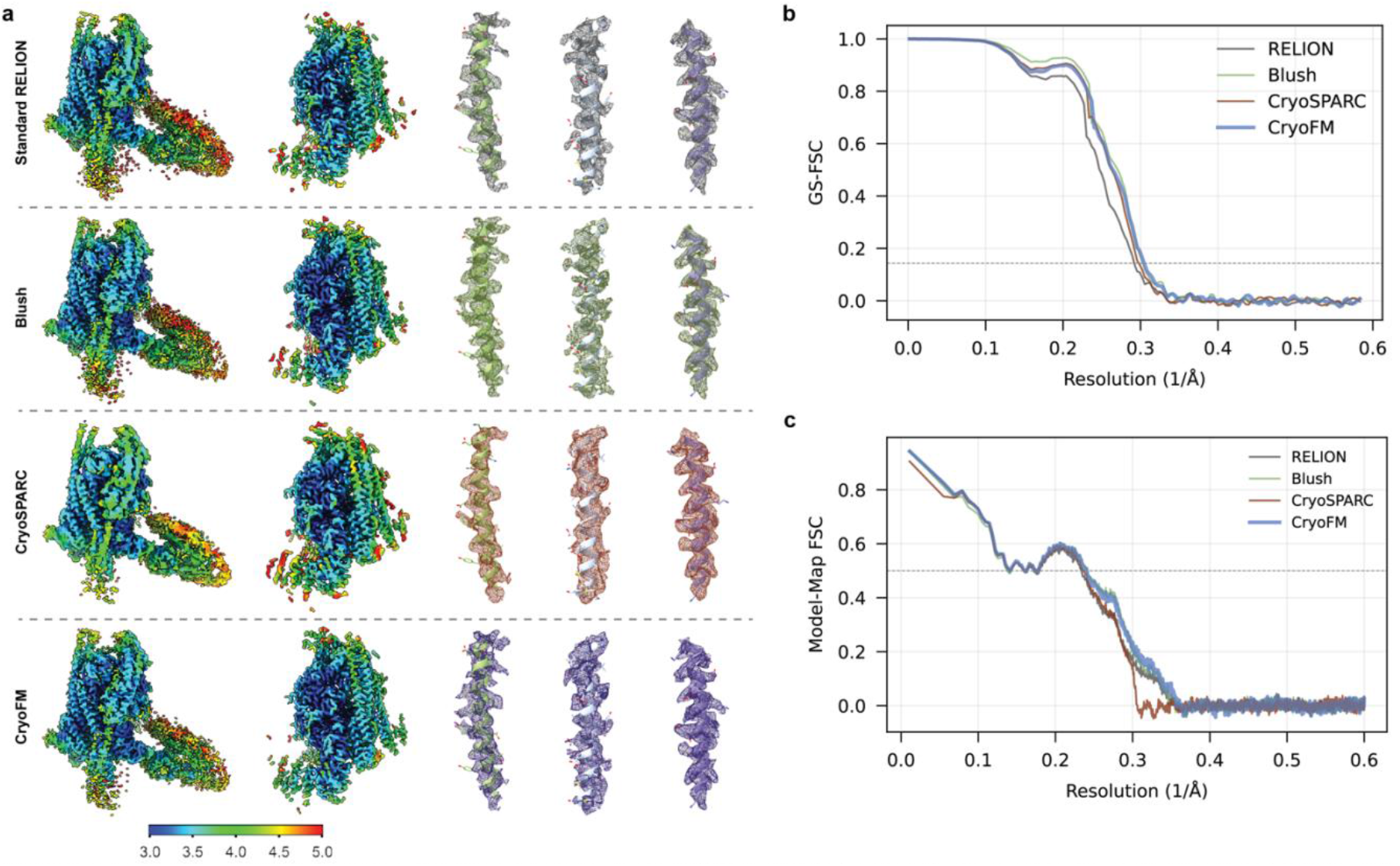
Comparison of refinement methods on EMPIAR-11910. **(a)** Density maps reconstructed using standard RELION, Blush, cryoSPARC, and cryoFM, shown with local resolution coloring (left) and representative zoom-in regions displayed with docked atomic models (right), **(b)** Half-map FSC curves and **(c)** model-map FSC curves of the four methods.

**Extended Data Figure 16.**
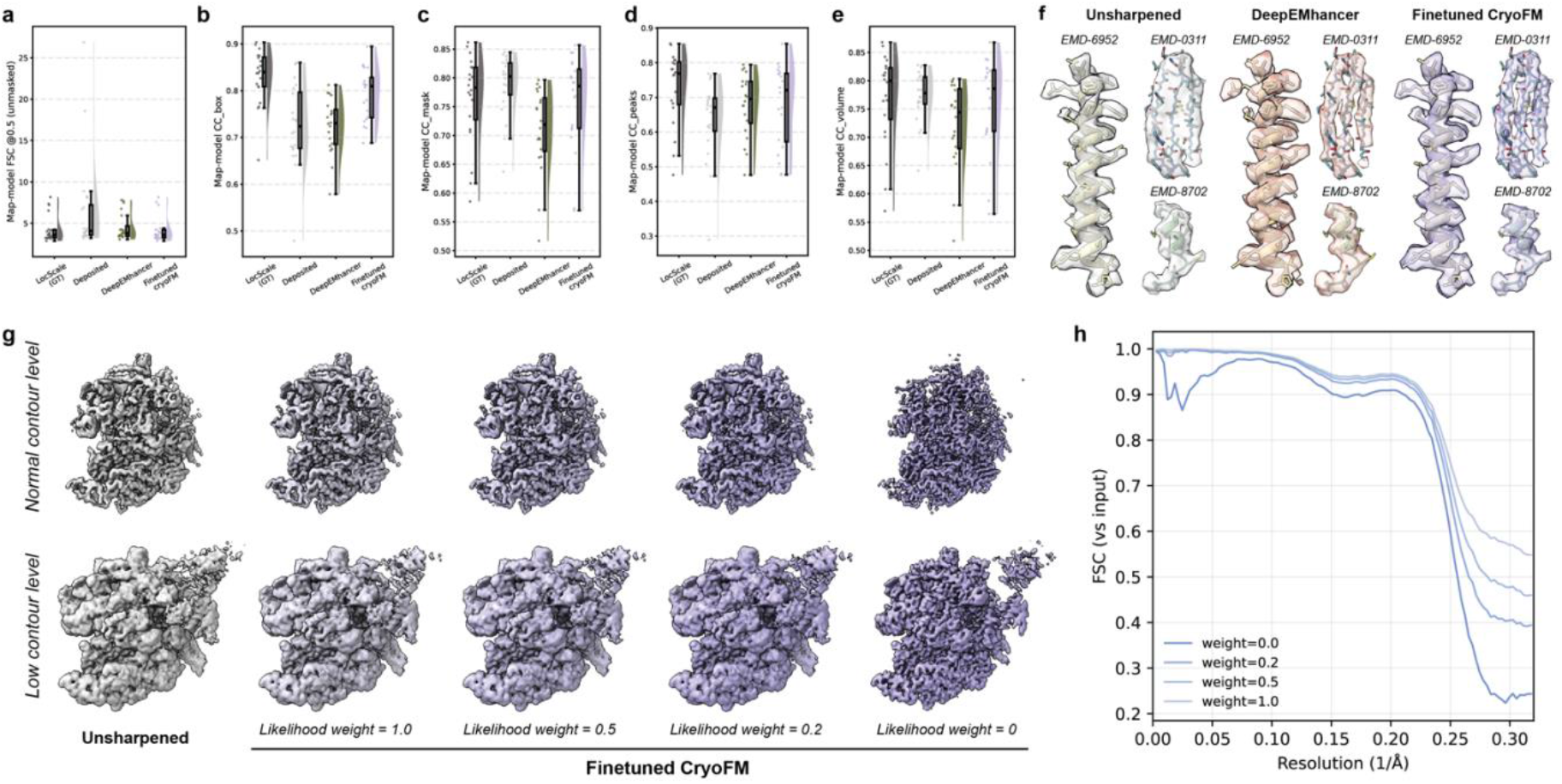
DeepEMhancer-style density post-processing with finetuned cryoFM and likelihood-weight control, **(a-e)** Quantitative benchmark comparisons of post-processed maps on the test set. **(f)** Representative density visualizations showing qualitative differences in local map features among post-processing methods, **(g)** CryoFM post-processed maps (EMD-0043) at normal and low contour levels under different likelihood weights, showing a continuous transition from prior-dominated, DeepEMhancer-style sharpening to likelihood-dominated enforcement of data consistency, **(h)** FSC curves calculated against the input density corresponding to the likelihood-weight titration.

**Extended Data Figure 17.**
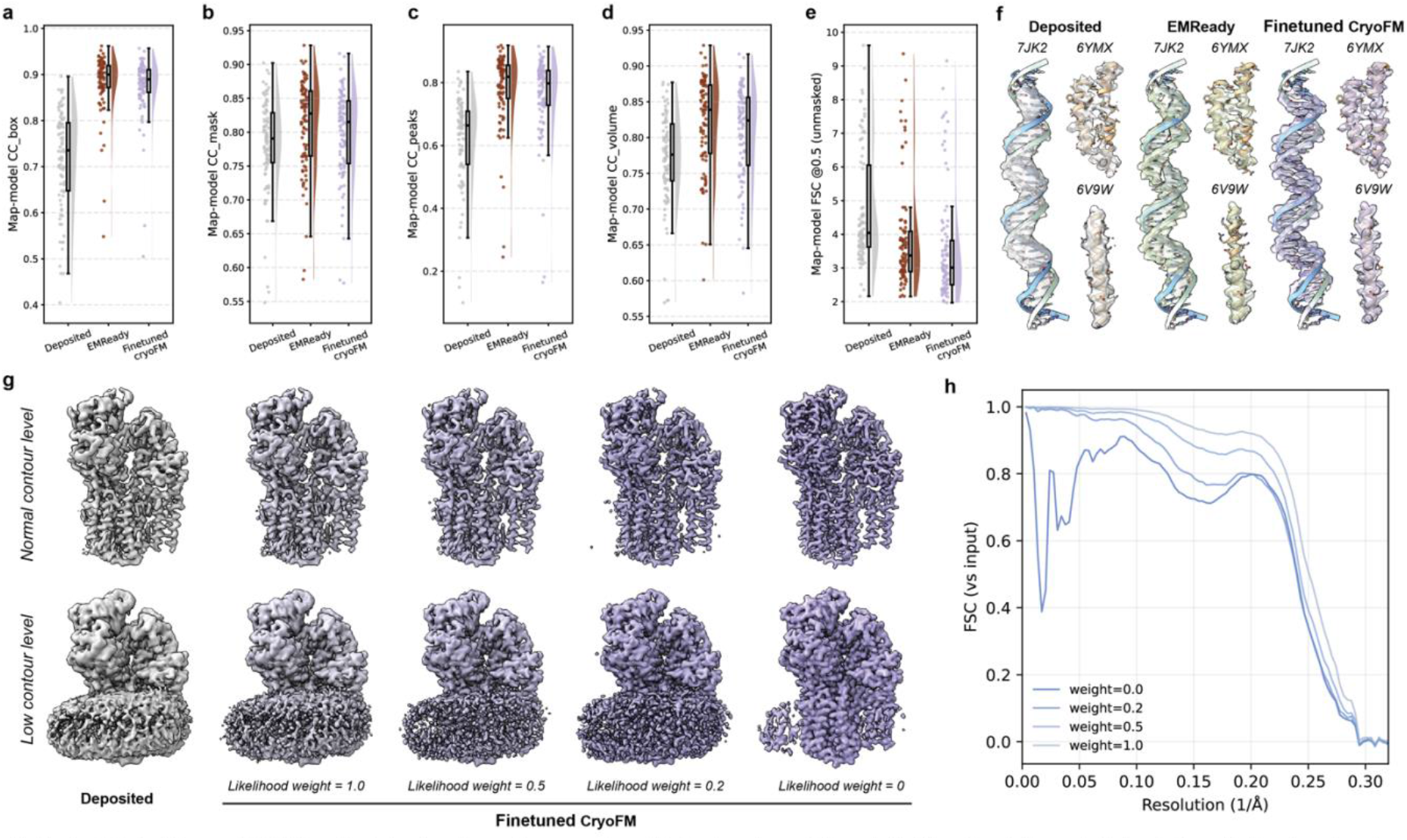
EMReady-style density post-processing with finetuned cryoFM and likelihood-weight control, **(a-e)** Quantitative benchmark comparisons of post-processed maps on the test set. **(f)** Representative density visualizations showing qualitative differences in local map features among post-processing methods, **(g)** CryoFM post-processed maps (EMD-22260) at normal and low contour levels under different likelihood weights, showing a continuous transition from prior-dominated, EMReady-style modification to likelihood-dominated enforcement of data consistency, **(h)** FSC curves calculated against the input density corresponding to the likelihood-weight titration.

## Supplementary Information

**Supplementary Table 1.** Summary of the notations.

| Symbol | Description | Space |
| --- | --- | --- |
| $D$ | Side length of the density map. | $\mathbb{N}$ |
| $N$ | Dimensionality of data space, $N = D^3$. | $\mathbb{N}$ |
| $M$ | Dimensionality of observation space, $M \leq N$. | $\mathbb{N}$ |
| $\mathbf{x}_0$ | Ground truth sample from the data distribution, $t = 0$. | $\mathbb{R}^N$ |
| $\mathbf{x}_t$ | Noisy state at time step $t$. | $\mathbb{R}^N$ |
| $\mathbf{v}_t$ | Velocity field of the flow-matching dynamics at time $t$. | $\mathbb{R}^N$ |
| $\mathbf{y}$ | Observed data used for conditioning. | $\mathbb{R}^M$ |
| $\mathbf{m}$ | Elementwise binary or soft mask applied to $\mathbf{x}$ or $\mathbf{y}$. | $\mathbb{R}^N$ or $\mathbb{R}^M$ |
| $\mathbf{r}$ | Coordinate in real space. | $\mathbb{R}^3$ |
| $\mathbf{k}$ | Coordinate in Fourier space. | $\mathbb{R}^3$ |
| $\mathcal{A}$ | Forward measurement operator, $\mathbf{y} = \mathcal{A}\mathbf{x} + \epsilon$. | $\mathbb{R}^N \rightarrow \mathbb{R}^M$ |
| $\mathcal{F}$ | Fourier transform operator. | $\mathbb{R}^N \rightarrow \mathbb{C}^N$ |
| $\mathcal{W}$ | Wavelet transform operator (with $B$ wavelet subbands). | $\mathbb{R}^N \rightarrow \mathbb{R}^{B \times N}$ |

### A Diffusion and flow matching

Diffusion models are a class of generative models that represent complex data distributions through a progressive noising and denoising process. In scientific modeling, this evolution is commonly formulated as score-based stochastic dynamics, yielding a natural stochastic differential equation (SDE) description of generation^1,2^. After training, new samples can be generated by simulating a reverse-time dynamics that starts from a simple reference distribution, such as a standard Gaussian, and gradually transforms it toward the target data distribution. Flow matching provides a closely related but deterministic alternative to diffusion-based generation. Rather than relying on stochastic reverse-time sampling, flow matching formulates generation as an ordinary differential equation (ODE) that deterministically transports samples from a simple base distribution to the data distribution^3^.

Diffusion- and flow-based generative models now play a central role in modern AI applications for structural biology and protein design, as they enable expressive modeling of rich, multimodal distributions in high-dimensional continuous spaces. Two aspects are particularly relevant in practice.

#### (i) Generative models for structure ensembles

Many problems require sampling diverse yet plausible structures that satisfy geometric, biochemical, or experimental constraints. Diffusion and flow-based models address this need by learning distributions over protein structures and complexes, enabling controllable generation under specified constraints. Representative examples include programmable generative models such as Chroma, which support controlled generation of protein structures and complexes^4^, and RFdiffusion, which enables conditional generation of scaffolds and binders under structural and functional constraints^5,6^. Joint sequence-structure models, such as ProteinGenerator, further extend this paradigm by learning multimodal distributions over both sequence and structure^7^, while all-atom generative frameworks such as Protpardelle capture conformational variability at atomic resolution^8^.

#### (ii) Conditional generation

Generative models can naturally incorporate partial observations through conditioning, enabling principled generation, completion and modification. This framework has been formalized for general inverse problems within score-based and diffusion models^9,10^, and has been applied in protein design through programmable conditional generation in Chroma^4^ and guided diffusion approaches such as RFdiffusion^5^.

Importantly, in these applications, generative models are not used merely as structure generators. Instead, they function as flexible probabilistic priors over high-dimensional scientific objects, enabling principled integration with conditions and/or experimental constraints. In this work, we adopt flow matching as the foundation-model framework because it provides a simple and direct formulation of generative modeling. By defining generation through a deterministic continuous-time ODE, flow matching enables stable sampling, straightforward incorporation of constraints and a unified backbone for unconditional generation, conditional modification, and task-specific adaptation.

### B Foundation models

Foundation models aim to capture broad, reusable knowledge from large and diverse datasets, enabling adaptation across many downstream applications with minimal task-specific tuning. Such models have transformed many scientific domains, particularly protein structure prediction and design. In computational biology, examples include AlphaFold3^11^, RFDiffusion^5^, and Chroma^4^, which use generative diffusion frameworks to learn distributions of atomic coordinates or protein backbones, as well as ESM-3^12^, which extends language-model pretraining to sequence generation. These models demonstrate that when trained at scale, a single generative prior can be adapted to many structure-related problems without retraining from scratch.

Conceptually, cryoFM extends the foundation model paradigm into the density domain of structural biology. Whereas AlphaFold3 and RFDiffusion build generative priors in atomic coordinate space, cryoFM builds them in density space, directly aligned with the experimental observables of cryo-EM and cryo-ET. This complementary perspective makes it applicable to experimental data processing and interpretation and map improvement. In this sense, cryoFM functions as a foundation model for cryo-EM, encapsulating learned structural priors that can generalize across various tasks.

### C Software and data availability

The source code used in this study is openly available at https://github.com/ByteDance-Seed/cryofm. The lists of datasets used for training and evaluation are publicly available at Zenodo under the DOI: https://doi.org/10.5281/zenodo.18013604. The pretrained and finetuned model weights are available at https://huggingface.co/ByteDance-Seed/cryofm-v2. A detailed user guide can be found at https://bytedance-seed.github.io/cryofm/docs. This paper is for research purposes only and not integrated into ByteDance technologies.

### D Additional information of posterior sampling

Posterior sampling methods such as diffusion- and flow-based formulations have recently emerged as powerful frameworks for addressing general inverse problems, where the goal is to recover an underlying signal from a corrupted or partial observation defined by a forward operator. These problems encompass a broad range of imaging and signal restoration tasks, including super-resolution, inpainting, and deblurring, all of which can be unified under the probabilistic relationship between the true sample, its measurement, and the observation model.

#### D.1 Problem setup and Bayes formulation

Let **x**_0_ ∈ ℝ^*N*^ denote a clean cryo-EM density and **y** ∈ ℝ^*M*^ an observation obtained by a forward operator 𝒜: ℝ^*N*^ → ℝ^*M*^ that models experimental degradations. We consider the time-indexed latent **x**_*t*_ produced by a pretrained generative model. By Bayes’ rule the posterior at time *t* satisfies

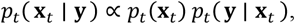

and the corresponding posterior score factorizes as

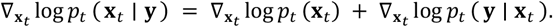

The first term is the prior score given by the pretrained model; the second is the likelihood score that enforces data consistency under 𝒜.

#### D.2 Diffusion posterior sampling (DPS): intuition and approximation

Direct evaluation of log *p*_*t*_ (**y** | **x**_*t*_) is generally intractable because it requires marginalizing over all clean signals consistent with **x**_*t*_. A practical approximation is to replace this marginal with a Laplace approximation around the model’s current estimate of the clean signal, 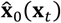. This yields:

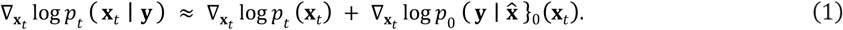

Assuming an observation model *p*_0_(**y** | **x**) with forward operator 𝒜,

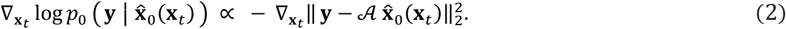

Combining (1)-(2) gives a working form of the posterior score used in diffusion posterior sampling:

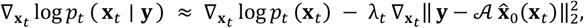

where λ_*t*_ absorbs proportionality constants and serves as a weight for the likelihood term.

#### D.3 Flow posterior sampling (FPS)

In a flow matching model, the prior distribution is parameterized by a time-dependent vector field **v**_Θ_(*t*, **x**_*t*_) that transports samples from noise to data. It is related to the prior score function through

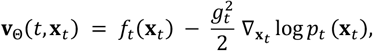

where *f*_*t*_ is a drift term and *g*_*t*_ is a diffusion scale.

The posterior (conditional) vector field follows from the decomposition of the posterior score:

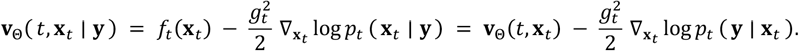

Under the rectified flow parameterization, 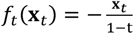 and 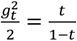, we have

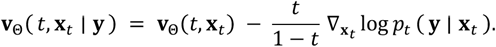

Using the Laplace approximation for the likelihood term yields a practical posterior flow:

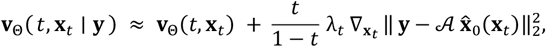

where λ_*t*_ controls the relative strength of the likelihood. In cryoFM, 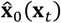 is the current estimate of the clean signal obtained from the flow state, *e*.*g*., 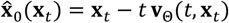.

In practice we use a stable ODE-style integrator and normalize the likelihood gradient to mitigate exploding updates. The factor 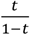 is clipped via *λ*_*max*_ avoid excessive corrections near *t* → 1 . FPS requires no retraining and task specificity is expressed through the likelihood *p*(**y** | **x**).

##### Algorithm 1

Flow Posterior Sampling (FPS)

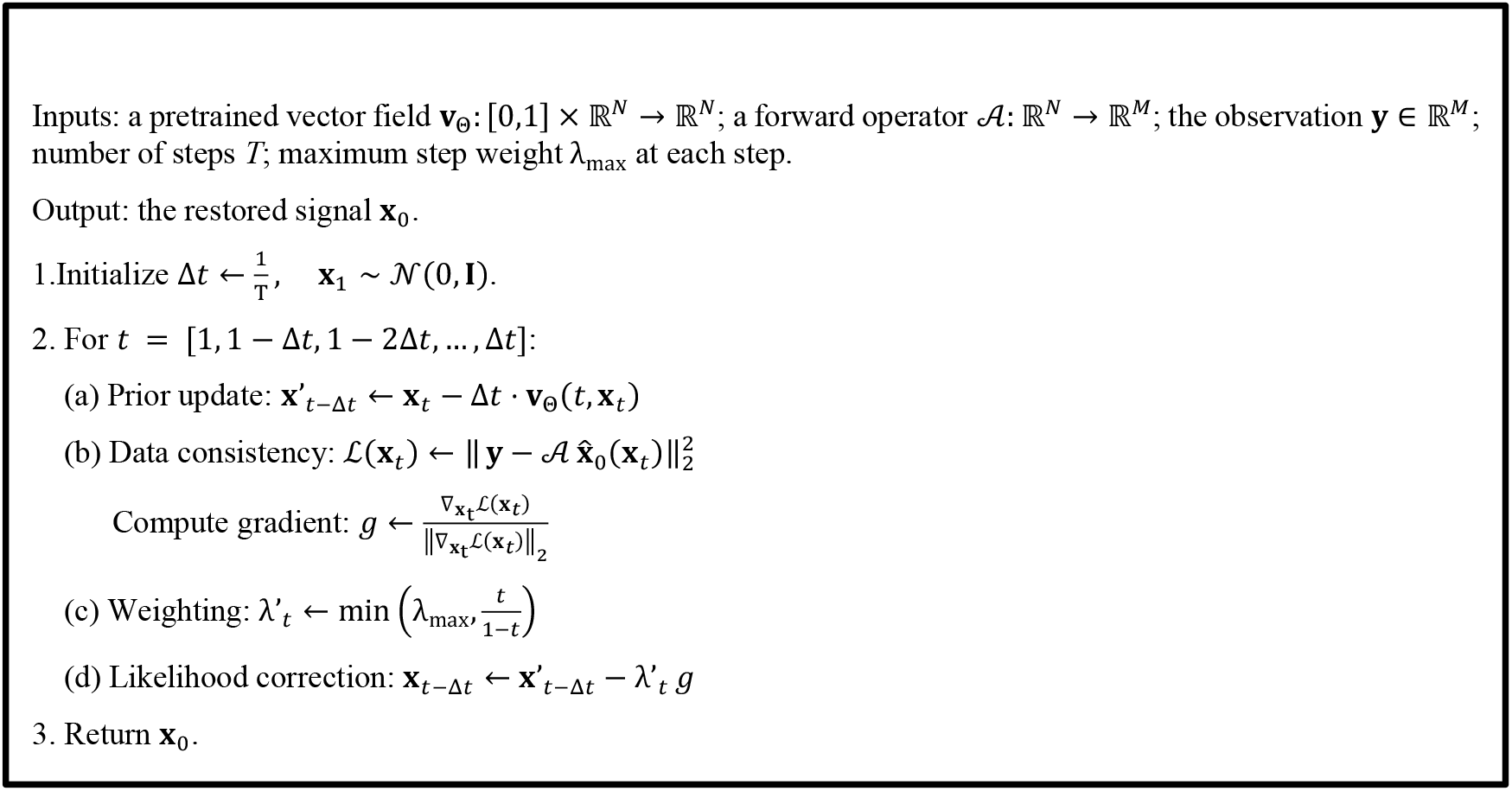

### E Derivations of forward operators

In the context of flow posterior sampling, the forward operator defines the likelihood term that links the clean signal to its observed measurement. Explicit derivation of these operators provides a physically interpretable bridge between the learned generative prior and the observed data. The parameters of these operators are derived from the empirical observations, typically the half maps obtained from independent reconstructions, or in some cases, the particle pose distribution estimated in refinement. For each task, we first define the analytical form of the forward operator that reflects the assumed degradation, and then infer its parameters from the observations. This formulation ensures that the forward operators used in posterior sampling are physically interpretable, task-specific, and quantitatively consistent with the actual experimental conditions.

#### E.1 Isotropic spectral noise variance

To estimate the noise power spectrum, we rely on two independent half maps, **y**_1_and **y**_2_, reconstructed from disjoint particle subsets. We denote by **X** and **Y**_*i*_ the Fourier transforms of the underlying signal **x** and the reconstructed maps **y**_*i*_, respectively. In Fourier space, we assume

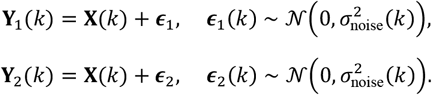

The gold-standard Fourier shell correlation (GS-FSC) between these two maps is:

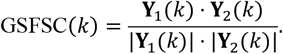

From GS-FSC, the shell-wise signal-to-noise ratio (SNR) is derived as^13^:

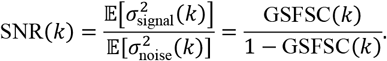

Since the expectation of the radial power spectrum decomposes into signal and noise contributions,

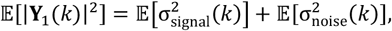

we can solve for the expected noise power for **Y**_1_:

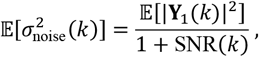

Alternatively, this is equivalent to:

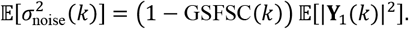

This provides a principled way to estimate frequency-dependent noise variances directly from half maps, yielding the spectral noise model used in posterior sampling.

#### E.2 Anisotropic spectral noise variance

To quantify the directional sampling distribution of particles in Fourier space, we construct a Fourier mask **m**^ori^ from the set of particle orientations used during reconstruction. Specifically, each particle’s projection direction is backprojected into a 3D Fourier grid, and the resulting contributions are accumulated over all particles. The value of **m**^ori^(**k**) at voxel coordinate **k** thus reflects how frequently that spatial frequency is sampled in the dataset, that regions near preferred orientations receiving higher weights and poorly sampled directions lower ones.

Normalizing **m**^ori^(**k**) it within each shell gives

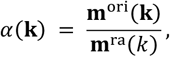

where **m**^ra^(*k*) = **m**^ori^(**k**)_|**k**|=*k*_, which satisfies ∑_|**k**|=*k*_ *α*(**k**) = 1. We then preserve the shell-average noise power while redistributing it directionally by setting:

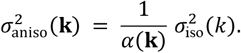

This yields the anisotropic forward operator above and matches the implementation that weights likelihoods by the inverse of the direction-specific noise variance.

#### E.3 Real space local heterogenous noise variance

To characterize spatially varying noise while retaining frequency specificity, we employ a 3D Shannon wavelet decomposition^14^. Unlike conventional Fourier-based or purely local filtering approaches, a wavelet representation provides a joint description of signal variations in both space and frequency, allowing local noise statistics to be analyzed without losing spectral information. Among available wavelet bases, the Shannon wavelet offers perfect frequency partitioning and energy-preserving band separation. Each sub-band corresponds to a non-overlapping radial frequency interval, and the inverse transform of an ideal band-limited component yields a smoothly varying real-space field without overlap between bands. This construction allows per-band local variance estimation to be performed directly in real space while maintaining a clear correspondence to specific frequency ranges.

Specifically, the Shannon basis partitions the Fourier domain into a set of non-overlapping, ideal band-pass filters, each defined by a binary radial mask 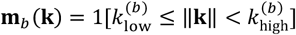, *b* = 1, …, *B*. The wavelet transform 𝒲 is implemented using ideal Shannon band-pass filters in Fourier space, followed by inverse Fourier transform to obtain real-valued responses. Formally, for a 3D density **x**,

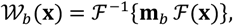

where **m**_*b*_ denotes the binary mask selecting frequencies within the *b* -th frequency band, and the multiplication **m**_*b*_ ℱ(**x**) is understood as pointwise multiplication.

The resulting set of coefficients 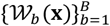 provides a localized multiscale representation of both signal and noise, which can be used to estimate spatially varying noise power and incorporated into posterior sampling through frequency band-wise and voxel-wise weighting.

Given two half maps **y**_1_ and **y**_2_ with independent noise of equal variance, we apply the 3D Shannon wavelet transform 𝒲 to both, yielding band-wise coefficients:

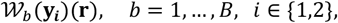

where *b* indexes the radial frequency bands and **r** denotes the voxel location in real space.

Under the standard half map assumptions (identical underlying signal, uncorrelated zero-mean noise with equal variance), the expected squared difference between the two half maps equals twice the local noise variance:

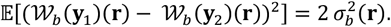

Hence, an unbiased estimator of the local, band-resolved variance is:

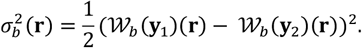

The estimator 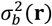 provides a per-voxel, per-frequency measure of spatially varying noise power, effectively combining real-space adaptivity and spectral resolution. Because the Shannon basis uses ideal band-pass filters, the resulting sub-bands partition the total signal energy without overlap, allowing simple and interpretable aggregation across frequencies. In practice, the number of bands *B* determines the trade-off between frequency precision and spatial smoothness.

### F Ablation on posterior sampling under the synthetic missing-cone setting

To assess the robustness of the proposed posterior sampling procedure in a controlled inverse problem setting, we consider a synthetic missing-cone experiment that mimics the effect of preferred particle orientations in cryo-EM reconstruction. This setup is similar to the synthetic experiment above on orientation-induced anisotropy, while allowing precise control over the severity of information loss.

Starting from a ground-truth density map **x**, we generate synthetic observations by removing Fourier coefficients within a symmetric conical region centered around the missing directions. The cone is parameterized by its half-angle *β*, with larger *β* corresponding to more severe loss of directional information. The observed volume is obtained by inverse Fourier transforming the masked spectrum, while coefficients inside the missing cone are set to 0. Posterior samples are then drawn using the unconditional cryoFM model, enforcing exact consistency with the observed Fourier coefficients and iteratively completing the missing region.

We evaluate reconstruction quality using Fourier shell correlation (FSC), reported both over the full Fourier domain (“whole map”) and restricted to the missing-cone region (“missing part”). The latter directly reflects the model’s ability to infer information that is not present in the observation. As a reference, we also report FSC between the synthetic observation and the ground truth, for which the missing-part FSC is identically zero by construction.

Supplementary Table 2 summarizes the effect of the number of posterior sampling steps for cone half-angles *β* = 30°, 45°, 60°. Two consistent trends emerge. First, increasing the number of sampling steps from a single iteration to a small number (approximately ten) leads to a clear improvement in both whole-map and missing-part FSC across all tested angles, suggesting that only a limited amount of posterior refinement is required to obtain stable reconstructions in this setting. Second, further increasing the number of sampling steps does not yield systematic gains. For larger missing cones (*β* ≥ 60°), longer sampling chains are associated with a gradual decrease in missing-part FSC and a mild degradation in whole-map FSC. This behavior reflects the increasing ill-posedness of the inverse problem as the missing cone widens: As the inverse problem becomes increasingly underdetermined in the missing Fourier region, additional sampling primarily explores variability rather than converging to a more accurate completion. As a result, excessively long posterior sampling can amplify uncertainty in the missing region without improving global fidelity.

**Supplementary Table 2.** Ablation of posterior sampling steps under a synthetic missing-cone setting. Average Fourier shell correlation (FSC) is reported for reconstructions obtained with different numbers of cryoFM posterior sampling steps, for cone half-angles *β* = 30°, 45°, 60°. Results are shown for the whole map and for the missing-cone region (“missing part”). The “Observation vs. GT” row reports FSC between the masked observation and the ground truth, for which the missing-part FSC is zero by construction.

| | $\beta = 30^\circ$ | | $\beta = 45^\circ$ | | $\beta = 60^\circ$ | |
| --- | --- | --- | --- | --- | --- | --- |
|  | Whole map | Missing part | Whole map | Missing part | Whole map | Missing part |
| Observation vs. GT | 0.9230 | 0 | 0.8380 | 0 | 0.7142 | 0 |
| CryoFM (1 step) | 0.9253 | 0.0421 | 0.8411 | 0.0467 | 0.7169 | 0.0425 |
| CryoFM (10 step) | 0.9559 | 0.6565 | 0.8727 | 0.4608 | 0.7386 | 0.2922 |
| CryoFM (100 step) | 0.9574 | 0.6423 | 0.8631 | 0.4157 | 0.7077 | 0.2335 |
| CryoFM (200 step) | 0.9590 | 0.6552 | 0.8638 | 0.4203 | 0.6935 | 0.2097 |
| CryoFM (500 step) | 0.9534 | 0.6255 | 0.8540 | 0.3848 | 0.6939 | 0.2131 |
| CryoFM (1000 step) | 0.9547 | 0.6332 | 0.8598 | 0.4069 | 0.6880 | 0.2012 |

### G Discussion on comparison with other methods

Density restoration in cryo-EM encompasses two conceptually related yet practically distinct objectives. During 3D reconstruction, restoration or denoising functions as a regularization mechanism within the iterative refinement loop, aiming to improve convergence stability, prevent overfitting to noise, and avoid suboptimal local minima. In contrast, density modification is typically applied after reconstruction has converged, with the goal of correcting systematic errors and enhancing the interpretability of the final map. Despite these differences, restoration methods in both contexts share the overarching goal of improving map quality. Broadly, existing approaches can be categorized into analytical and deep learning-based methods. Analytical techniques offer explicit interpretability but often depend on heuristic assumptions and handcrafted rules. Deep learning-based approaches, on the other hand, can be further divided into two classes: self-supervised methods, which train a new model on the fly using only the current dataset and its inherent noise characteristics (*e*.*g*., paired half maps from the same reconstruction), and pretrained models, which are trained on curated datasets of EMDB maps to learn generalizable transformations. In the following subsections, we discuss and compare cryoFM with representative methods from each of these categories.

#### G.1 Comparison with analytical methods

Analytical approaches to density restoration and modification in cryo-EM have a long history and remain highly interpretable due to their explicit mathematical formulations. Under the Bayesian framework, RELION models reconstruction as the maximization of a posterior distribution with a Gaussian prior on the density^15^. This prior effectively suppresses high-frequency components with low signal-to-noise ratios, acting as a global smoothness regularizer. Later developments, such as non-uniform refinement^16^ and SIDESPLITTER^17^, extended this idea by introducing local rather than global regularization, heuristically adapting the degree of smoothing across space to improve convergence and reconstruction quality. In the realm of density modification, analytical methods typically operate in Fourier space, employing frequency-dependent weighting or filtering schemes. Examples include Wiener-style filtering^18^, local resolution-guided amplitude scaling^19^, and various Fourier shell-based corrections. Some approaches require additional external information (*e*.*g*., an atomic model) to provide structure-dependent weighting or phase constraints^20^.

The key advantage of analytical methods lies in their interpretability and controllability: every operation can be explicitly expressed as a formula, allowing users to understand and directly tune how regularization or modification affects the map. However, their limitations stem from the same explicitness, that they rely on heuristic assumptions and handcrafted rules, and thus struggle to generalize or capture the complex patterns observed in real cryo-EM data.

Compared with analytical approaches, which rely on explicitly defined regularizers or handcrafted filters, cryoFM unifies physical interpretability and data-driven expressiveness. Through FPS, it retains the explicitness of analytical formulations via the likelihood term while drawing on the rich structural statistics learned by the generative prior. Instead of designing heuristic regularizers or inverting degradation models, cryoFM only requires specifying a physically grounded forward operator that describes the observation process, making it both principled and easier to model.

#### G.2 Comparison with self-supervised methods

Self-supervised methods such as M^21^ and spIsoNet^22^ are commonly incorporated into the iterative refinement stage of cryo-EM reconstruction. Their primary objective is to suppress noise while preserving structural signal, often implemented within a noise2noise learning framework^23^ that uses the two independent half maps as paired noisy observations of the same underlying density. By exploiting only the data at hand, these methods aim to improve the effective signal-to-noise ratio without introducing external biases.

Because cryo-EM datasets typically exhibit extremely low SNR, structural biologists are very cautious about introducing any external information into the refinement loop, as even small biases can distort the final structure. This caution has made self-supervised learning particularly appealing: it combines the representational power of deep neural networks with the perceived safety of using only internally consistent data from the same reconstruction.

Despite this seemingly advantage, self-supervised approaches are inherently constrained by their closed data scope. The absence of external training data means that these models cannot benefit from the rich structural variability present across macromolecules, and the two half maps alone may not provide sufficient statistical diversity for optimal learning. Furthermore, since each dataset requires its own model, these methods demand additional training effort and are difficult to scale. Even after training, the learned denoiser remains a black box, and it may still introduce artificial or hallucinated features.

CryoFM extends the self-supervised emphasis on safety and dataset specificity to a broader, more data-efficient framework. It incorporates global structural knowledge learned from large-scale data as a pretrained prior, while still allowing dataset-specific adaptation through the likelihood term. This eliminates the need for retraining on each dataset and enables controlled, interpretable restoration guided by both prior knowledge and current observations.

#### G.3 Comparison with supervised pretrained methods

Supervised pretrained models are widely used in density modification tasks. Representative examples include DeepEMhancer^24^ and EMReady^25^. These models are typically trained in a supervised manner on curated datasets, where each training pair consists of an input map and its postprocessed counterpart. The targets can vary from locally sharpened maps to simulated densities derived from atomic models or other processed modalities, allowing the model to learn a deterministic mapping from input to output representations.

Blush^26^ extends this supervised concept directly into the refinement process. It constructs training pairs by simulating common degradation effects on cryo-EM densities and trains a neural denoiser to replace the analytical regularizer in RELION. To mitigate potential bias, Blush applies a heuristic spectral trailing strategy that retains only frequencies where the gold standard FSC exceeds 0.143. This threshold provides a conservative safeguard for defining the resolution limit by retaining only statistically reliable frequencies, but correlations above the 0.143 threshold may still be overestimated.

Supervised pretrained models, when well trained, can achieve impressive denoising or restoration performance, while their deterministic inference makes them extremely fast during application. However, the models are largely black boxes, offering little interpretability or controllability, and their generalization ability is constrained by the construction of the training pairs, making them vulnerable to out-of-distribution artifacts when applied to unseen data.

Unlike supervised pretrained models that learn a fixed mapping from noisy to clean maps, cryoFM serves as a flexible generative prior within a unified Bayesian formulation. Its explicit FPS framework allows different restoration or enhancement tasks to be addressed simply by defining task-specific likelihoods, rather than constructing separate training pairs for each objective. When fine-tuned on the same curated datasets used by supervised methods such as DeepEMhancer or EMReady, cryoFM achieves superior performance by leveraging its broader pretraining and more expressive prior. A remaining limitation of cryoFM is computational efficiency, as its iterative sampling process is slower than the one-step inference used in deterministic pretrained models.

### H Discussion on the limitation of FSC as the evaluation metric

For more than a decade, the gold-standard Fourier shell correlation (GS-FSC) has been widely accepted as a reliable evaluation metric in cryo-EM^13^. It serves not only as a resolution estimator (through the 0.143 cutoff threshold), but also as a general measure of reconstruction quality. The underlying idea is elegant: by splitting the dataset into two halves and refining them independently, the correlated signal represents reproducible structural features, while the uncorrelated components are treated as noise. This design has long provided a principled way to assess overfitting and validate reconstruction fidelity.

However, as modern algorithms become increasingly sophisticated, the reliability of GS-FSC is being challenged. Many recent reconstruction frameworks introduce complex regularization or denoising steps at each iteration, and some even employing deep neural networks. While these methods improve map quality, they can also introduce shared biases between the two half reconstructions. When both halves share the same information-rich or biased regularizer, they may exhibit spurious correlations, even if the refinement procedures themselves remain nominally independent. Similar effects can also arise from implicit neural representation models such as NeRF, where the inductive biases of the network architecture can impose correlated structural patterns across the two half reconstructions^27,28^. CryoFM employs a Bayesian formulation with an informative but broadly distributed data-driven prior, learned from diverse macromolecular densities, and stochastic posterior sampling that helps reduce deterministic bias. Still, such measures only mitigate rather than eliminate the susceptibility of GS-FSC to inflation caused by shared priors or modeling assumptions.

Taken together, we believe that GS-FSC, and other half map-based metrics^29,30^ (*e*.*g*., 3D-FSC), are increasingly vulnerable to method-dependent artifacts. Independence of the half datasets alone is no longer sufficient to guarantee unbiased evaluation. When comparing methods, relying solely on GS-FSC-based metrics can be misleading; qualitative assessment of overall map quality is becoming more essential.

Beyond algorithmic bias, we found that FSC itself as a metric can also be artificially inflated by certain real-space numerical operations, leading to overestimated correlations even without any change in the underlying information. For example, in several experiments, we observed that applying a simple non-negativity clip *x* ← max(*x*, 0) to a reconstructed density can raise its FSC to the “ground truth” (GT). We speculate that two mechanisms may contribute to this effect. First, negative densities mainly arise as mathematical artifacts of Fourier-space operations rather than representing any physical signal. Truncating these values removes the negative tail of the distribution, effectively reducing variance in noise-dominated regions. This can artificially increase the spectral SNR and, consequently, the FSC. Second, single-particle reconstruction relies on the Fourier slice theorem. Non-uniform pose distributions or missing views introduce anisotropic sampling in Fourier space, leaving certain directions undersampled. The real-space truncation *x* ↦ *x*_+_: *x***1**_**x**>**0**_ acts as a multiplicative mask, which in Fourier space corresponds to a data-dependent convolution. This convolution redistributes energy across neighboring frequencies and smooths directional gaps, producing an apparent rise in FSC. Importantly, such increases do not reflect genuine information gain but rather the metric’s sensitivity to these regularization effects. In other words, real-space numerical operations can artificially elevate correlations between maps, even though these manipulations do not introduce any new or meaningful structural information.

### I More discussion of current challenges and future directions

The framework introduced in this work establishes a principled way to integrate learned generative priors with explicitly defined degradation models through Bayesian flow posterior sampling. Within this framework, several important challenges and opportunities remain. We believe that future developments can be broadly organized along three complementary directions: improving the prior model, refining likelihood and degradation operators, and advancing posterior sampling methodologies. Progress along all three axes will be critical for fully realizing the potential of generative inference in cryo-EM.

#### I.1 Improving generative priors for cryo-EM densities

An important direction for future work is the development of more expressive and uncertainty-aware generative priors. In the current formulation, cryoFM learns a distribution over high-quality cryo-EM densities, but uncertainty is not explicitly parameterized across spatial locations or frequency bands. This limited our training set to only the density maps with high resolution. In practice, however, uncertainty in cryo-EM reconstructions is strongly frequency dependent and is routinely estimated through metrics such as GS-FSC. A promising extension would be to incorporate frequency-dependent uncertainty estimates directly into the prior or its interaction with the likelihood. By explicitly quantifying uncertainty at different spatial frequencies, such a model could leverage a much broader range of experimental datasets, including those with lower resolution. In this setting, lower-confidence frequency components would contribute with appropriately reduced weight, ensuring that useful information is not discarded while preventing unreliable signal from dominating the inference. This would allow generative priors to be applied more broadly without sacrificing robustness.

Another limitation of the current prior lies in its ability to capture long-range, global structural information. Practical implementations are constrained by patch-based training and finite receptive fields, which may limit sensitivity to global shape, symmetry, or long-range correlations. Future prior models that better capture global context, either through hierarchical representations, autoregressive architectures, or explicit global conditioning, could further improve performance, particularly for large complexes or assemblies with extended structures.

#### I.2 Advancing posterior sampling and inference strategies

Posterior sampling is central to the framework proposed here, but its current instantiation leaves room for substantial improvement. One immediate challenge is computational efficiency. While flow-based posterior sampling is more efficient than many stochastic alternatives, sampling remains relatively slow for large-scale or iterative reconstruction tasks. A second major challenge is inference under partially known or fully unknown degradation operators. In many realistic settings, parameters of the forward operator are not fully known *a priori*. Extending posterior sampling to blind or semi-blind settings^31^, where parameters of the degradation operator are jointly inferred alongside the density, represents an important direction. More generally, posterior sampling methods that can accommodate degradation operators that are only partially specified, or defined implicitly through data-driven models, would greatly expand the scope of applications. Such extensions would allow the framework to move beyond hand-crafted likelihoods while retaining Bayesian interpretability.

#### I.3 Toward unified and more expressive and accurate degradation models

A third important direction concerns the design of more unified, expressive and statistically consistent degradation operators. In the current framework, different sources of degradation are handled through separate likelihood formulations. A more unified forward operator that captures multiple degradation mechanisms simultaneously would simplify inference and better reflect the true data-generating process of cryo-EM experiments. Beyond unification, an important opportunity lies in expanding the types of degradation operators considered. For example, operators analogous to particle polishing^32^ could be incorporated to model per-particle beam-induced motion or dose-dependent degradation in a principled probabilistic manner. Similarly, degradation operators that explicitly address flexible heterogeneity, either through continuous deformation models or low-dimensional latent representations, could enable joint inference of structure and conformational variability within the same Bayesian framework.

Accurate estimation of degradation parameters from experimental observations is also crucial. From a statistical perspective, current approaches often rely on frequency-space operators estimated globally from the full map, which are then applied locally along densities in real space. This mismatch may not be fully consistent and warrants further investigation. Developing degradation models that are statistically coherent across spatial and frequency domains remains an open challenge. Moreover, while current implementations primarily rely on half maps, additional sources of information could be exploited to improve parameter estimation and likelihood design. Another promising direction is to represent degradation operators themselves using pretrained neural networks. In scenarios where the forward process is difficult to specify analytically or only partially understood, data-driven operators learned from large datasets could serve as flexible approximations within the posterior sampling framework. Such neural operators could be combined with explicit physical constraints, offering a hybrid approach that balances interpretability and expressiveness.

Finally, deeper integration of the cryo-EM forward imaging process, particularly the 3D-to-2D projection, represents an important frontier. Incorporating more accurate projection models within the posterior sampling framework may require tighter coupling with expectation-maximization or stochastic gradient-based optimization^33^ used in refinement. Such integration could lead to more robust pose inference, faster convergence and improved stability in challenging datasets.

